# Loss of *SMARCB1* evokes targetable epigenetic vulnerabilities in Epithelioid Sarcoma

**DOI:** 10.1101/2024.09.18.613695

**Authors:** Jia Xiang Jin, Fabia Fuchslocher, Martha Carreno-Gonzalez, Felina Zahnow, A. Katharina Ceranski, Rainer Will, Dominic Helm, Felix Bestvater, Ana Banito, Roland Imle, Shunya Ohmura, Florencia Cidre-Aranaz, Thomas G. P. Grünewald

**Affiliations:** Hopp-Children’s Cancer Center (KiTZ), Heidelberg, Germany; Division of Translational Pediatric Sarcoma Research (B410), German Cancer Research Center (DKFZ), German Cancer Consortium (DKTK), Im Neuenheimer Feld 280, 69210 Heidelberg, Germany; Medical Faculty, Ruprecht-Karls-University, Im Neuenheimer Feld 672, 69120 Heidelberg, Germany; National Center for Tumor Diseases (NCT), NCT Heidelberg, a partnership between DKFZ and Heidelberg University Hospital, Germany; Core Facility Cellular Tools (W111), German Cancer Research Center (DKFZ), German Cancer Consortium (DKTK), Heidelberg, Germany; Core Facility Proteomics (W120), German Cancer Research Center (DKFZ), German Cancer Consortium (DKTK), Heidelberg, Germany; Light Microscopy Core Facility (W210), German Cancer Research Center (DKFZ), German Cancer Consortium (DKTK), Heidelberg, Germany; Soft-Tissue Sarcoma Junior Research Group, German Cancer Research Center (DKFZ), German Cancer Consortium (DKTK), Heidelberg, Germany; Faculty of Biosciences, Heidelberg University, Heidelberg, Germany. Division of Pediatric Surgery, Department of General, Visceral and Transplantation Surgery, University Hospital Heidelberg, Heidelberg, Germany; Institute of Pathology, Heidelberg University Hospital, Heidelberg, Germany

**Keywords:** Epithelioid sarcoma, SWI/SNF, SMARCB1, epigenetics, therapeutic vulnerability

## Abstract

Dysfunction of epigenetic modulators, such as the SWI/SNF complex, is a wide-spread but relatively ill-defined feature of a broad spectrum of cancer entities. Among SWI/SNF-mutant entities, *SMARCB1*-deficient cancers, such as the highly aggressive Epithelioid Sarcoma (EpS), are characterized by this genetic event in an otherwise rather silent mutational landscape. This renders EpS an ideal model to study how epigenetic reprogramming by a single mutation can contribute to tumorigenesis.

Hence, to characterize and compare the function of the *SMARCB1*-deficient, residual and the physiological SWI/SNF complex in cancer, we generated a panel of *SMARCB1* re-expressing EpS cell lines and employed a functional multi-omics approach. Here, we show that SWI/SNF holds canonical characteristics of both tumor-suppressors and proto-oncogenes due to its multi-faceted role in the regulation of the epigenome. Our data indicates that the loss of *SMARCB1* causes an overall loss of SWI/SNF chromatin affinity at *cis*-regulatory enhancer elements, inducing a preference for uncontrolled proliferation and cell cycle progression as opposed to development and differentiation. We further demonstrate that EpS cell lines depend on residual SWI/SNF action to maintain clonogenicity and proliferation. Consequently, EpS cell lines exhibit markedly increased sensitivity to pharmacological inhibition of the residual SWI/SNF when compared with SWI/SNF-proficient cancer entities.

Collectively, our results from the EpS model shed new light on how a single mutation can rewire the pleiotropic effects of an epigenetic master regulator and provide inroads for therapeutic intervention.

## INTRODUCTION

Epithelioid sarcoma (EpS) is a high-grade malignancy of unknown histogenesis first described in 1970 (ref. ^1,2^). It accounts for less than 1% of all adult sarcomas and 4–8% of pediatric soft tissue tumors outside the rhabdomyosarcoma-spectrum^3,4^. EpS consists of two main types: a more common and comparatively less aggressive distal (classical) type, mainly affecting the extremities (fingers, arms, and feet), as well as a less common but more aggressive proximal type mainly occurring in the trunk or upper extremities^2,5^. Despite multimodal therapies combining often mutilating surgery with radiochemotherapy, EpS features high rates of relapse and metastasis, resulting in unfavorable overall survival (5-year-survival rates of 60–75%^4^; 10-year survival rates of 42–55%^2^). In terms of targeted therapy, the U.S. Food and Drug Administration (FDA) has recently approved the enhancer of zeste homology 2 (EZH2) inhibitor tazemetostat for clinical use in EpS, which achieved transient objective responses in only a minority of patients (15%)^6^, possibly because of rapid development of drug-resistance^7^. Hence, a better understanding of the molecular pathomechanisms driving EpS is necessary to develop specific therapies that can reduce treatment-associated toxicity and provide durable objective responses in the majority of patients.

As a diagnostic hallmark, EpS is defined by a loss of nuclear Switch/Sucrose Non-Fermentable (SWI/SNF)-related, matrix associated, actin dependent regulator of chromatin, subfamily B, member 1 (*SMARCB1*) expression (alias BRG1/BRM-associated factor 47, BAF47; Integrase Interactor 1, INI-1; and human sucrose non-fermentable 5, hSNF5) which is caused by genomic deletions at the *SMARCB1* locus in most cases^8–10^. *SMARCB1* is a core subunit of the highly conserved mammalian SWI/SNF complex –a master regulator of chromatin remodeling and gene expression^11^. In fact, ∼20% of cancers exhibit tumorigenic perturbations to the SWI/SNF complex^11^. Crystallographic structure analyses show that *SMARCB1* acts as a molecular clamp coordinating SWI/SNF genomic affinity and site specificity alongside either of the two mutually exclusive core ATPases (BRM/SWI2-related gene 1 (BRG1) and Brahma (BRM)) by interaction with histone acidic patches^12^. The SWI/SNF complex in mammals occurs in three distinct assemblies, each characterized by a different composition of the subunits and chromatin affinity: canonical, polybromo-associated, and non-canonical SWI/SNF (cBAF, PBAF, ncBAF, respectively) with the former two incorporating *SMARCB1* as a core subunit^12,13^. The characteristic loss of *SMARCB1* as the putative oncogenic driver of EpS can also be found across a wide range of other malignancies, such as hematological malignancies, colorectal, ovarian, hepatocellular, squamous cell, breast and lung cancer^11^. Mechanistically, its role as a tumor supressor^14^ as well as the oncogenic properties gained by the residual SWI/SNF complex (rSWI/SNF) after loss of *SMARCB1* function^15^ have been suggested to drive tumorigenesis. Investigations into distinct entities exhibiting similar genomic alterations as EpS, such as malignant rhabdoid tumor (MRT), have shown a potentially context-dependent role in the maintenance^16^ and determination of cellular identity^17^ due to the functional relation of SWI/SNF with cell cycle progression^18^ and developmental pathways^19,20^. Perturbations to SWI/SNF assembly and subunits appear to greatly affect interactions with a range of transcription factors – most noticeable those of the AP-1 family^21^. Furthermore, nucleosome affinity and thus also downstream gene regulation appears to be dependent on SWI/SNF assembly^22^. Another proposed mechanism for the epigenetic regulation of this broad spectrum of downstream pathways lies in the antagonism (or lack thereof) of the polycomb repressive complex 2 (PRC2; containing the therapeutic target of tazemetostat (EZH2) as its catalytic subunit), a master repressor of gene expression, and the interaction with distal enhancer sites^23^.

Prior investigations showed that constitutive re-expression of *SMARCB1* had a strong anti-proliferative effect on MRT cell lines^23,24^. However, preliminary studies for EpS cell lines were less consistent in that respect, which may be due to the constitutive re-expression systems employed inducing adaptive cellular processes potentially obscuring the effects of immediate *SMARCB1* re-expression^23,24^. In MRT, it was shown that the residual SWI/SNF complex coordinates the epigenetic modulation of super-enhancers essential for tumor survival^25^. In keeping with these results, studies in MRT, acute myeloid leukemia, and prostate cancer models, all with proven dependency on the (residual) SWI/SNF complex, have shown that inhibition of the aberrant SWI/SNF-complex by small molecules or proteolysis-targeting chimeras (PROTACs) could offer new therapeutic possibilities to exploit the potential of synthetic lethality with low off-target toxicity^26–29^.

In this study, we take a functional genomics approach to characterize the role of *SMARCB1* in EpS by utilizing a conditional SMARCB1 re-expression system in several EpS cell lines. Thereby, we demonstrate a prominent *SMARCB1*-dependency of EpS on complex phenotypes affecting cellular identity, control of differentiation, cell cycle progression, proliferation, and apoptosis as well as epithelial-to-mesenchymal transition (EMT) programs. Furthermore, through multi-omics analyses we provide mechanistic evidence for pharmacologic inhibition of the residual SWI/SNF-complex as a therapeutic strategy in SWI/SNF-deficient entities such as EpS.

## RESULTS

### *SMARCB1* deletion promotes clonogenic growth and tumorigenicity of EpS

Loss of *SMARCB1* expression is considered a diagnostic hallmark of EpS, yet definitive evidence for its functional contribution to EpS malignancy is rather scarce, merely based on a limited set of cell line models, and not yet conclusively established^23,24^. To thoroughly investigate the role of *SMARCB1*, we generated isogenic models with doxycycline-(DOX)-inducible *SMARCB1* mRNA and respective empty vector controls by lentiviral transduction in a broad panel of six EpS cell lines. The parental cell lines (FU-EPS-1; HS-ES-1, -2M, -2R; NEPS; VA-ES-BJ) all exhibit homozygous *SMARCB1* deletion and represent both proximal- and distal-type EpS^2^. As shown in **Fig. 1a**, all cell lines exhibited prominent *SMARCB1* re-expression upon DOX treatment as verified by western blot. The DOX concentrations were then adjusted so that levels of *SMARCB1* re-expression determined using SYBR/TaqMan qPCR were similar to those observed in *SMARCB1* proficient Ewing sarcoma (EwS) cell line models. EwS cell lines exhibit no genetic alterations at the *SMARCB1* locus and were thus used as an estimate for physiological levels that appear otherwise similarly conserved across many non-cancerous cell lines^30,31^ (**Supp. Fig. 1a**). Additionally, western blots of cytosolic and nuclear fractions of EpS cells demonstrated that re-expressed SMARCB1 translocated to the nucleus while co-immunoprecipitation of the nuclear fraction against BRG1 – one of the two core ATPases present in all known SWI/SNF assemblies – showed that nuclear SMARCB1 readily re-incorporated into the SWI/SNF complex (**Fig. 1b**).

**Fig. 1:**
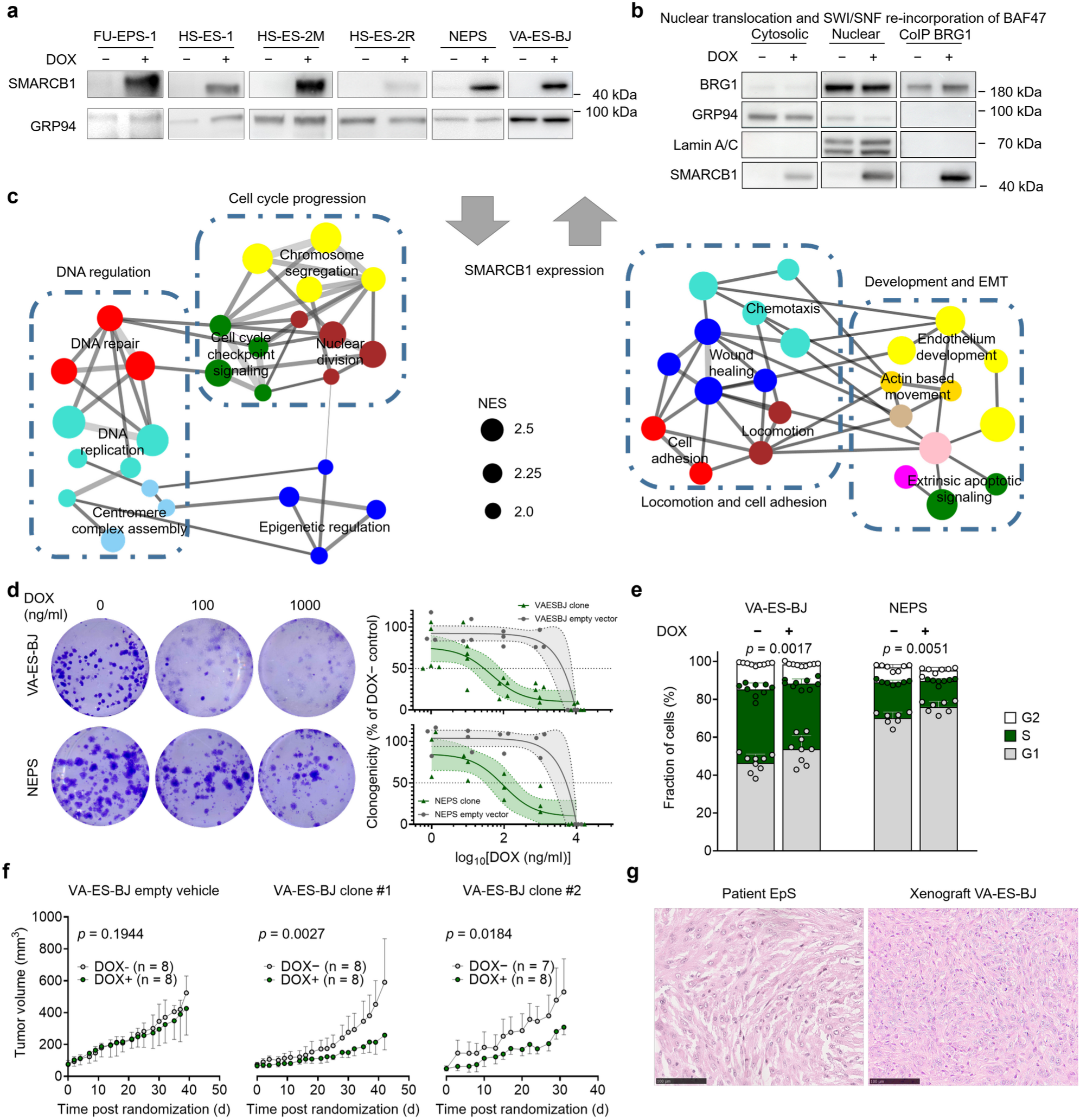
(a) Western Blot of clonal EpS cell line re-expressing SMARCB1. (b) Western blot of cytosolic, nuclear and BRG1-coIP‘d cell lysates against BRG1, GRP94 (cytosolic), Lamin A/C (nuclear) and SMARCB1 in VA-ES-BJ. (c) GSEA-based network analysis of up-/down-regulated gene sets upon SMARCB1 re-expression from shared regulated genes in VA-ES-BJ and NEPS. (d) DOX dosage-dependent gradual loss of clonogenicity in VA-ES-BJ and NEPS: Exemplary wells and fitted clonogenicity response curves. (e) Delayed G1/S-phase cell cycle progression upon SMARCB1 re-expression in VA-ES-BJ and NEPS. (f) *In vivo* tumor growth curves of VA-ES-BJ s.c. xenografts with empty vehicle or clonal SMARCB1 re-expression. (g) Comparison of HE-stained patient and xenograft tumour tissue.

As a first step in investigating the potential role of *SMARCB1* loss in EpS, we subjected two representative cell lines (distal type NEPS, proximal type VA-ES-BJ) with/without *SMARCB1* re-expression and respective controls to transcriptome profiling using Affymetrix Clariom D microarrays. In agreement with the putative oncogenic role of *SMARCB1* loss, comparative Weighted Gene Correlation Network Analysis (WGCNA) between both conditions based on gene set enrichment analysis (GSEA) showed significantly enriched gene sets mainly implicated in DNA repair and epigenetic regulation (downregulated) as well as several cellular differentiation and developmental pathways (upregulated), suggesting a role in tumor maintenance for the loss of *SMARCB1* (**Fig. 1c**).

To verify this prediction, we carried out various functional *in vitro* and *in vivo* experiments. We first conducted colony formation assays (CFAs) to investigate clonogenic growth as an indicator of oncogenic transformation^32^. Strikingly, both cell lines exhibited a dose-dependent reduction of clonogenicity upon *SMARCB1* re-expression (**Fig. 1d**). In agreement with these CFA and WGCNA results, flow cytometric cell cycle analysis using propidium iodide (PI) showed a delayed transition through G1/S-phase upon *SMARCB1* re-expression (**Fig. 1e**).

To confirm the tumor-suppressive function of *SMARCB1* in EpS, we conducted orthotopic, subcutaneous (s.c.) xenotransplantation experiments using VA-ES-BJ as a representative cell line in immunocompromised *Nod/Scid/gamma* (NSG) mice (**Fig. 1f**). Histological analyses confirmed that the xenografts recapitulated the typical morphology of EpS (**Fig. 1g**). Once tumors were palpable, *SMARCB1* was induced by DOX supplementation in drinking water, which led to a significant decrease of tumor growth as compared to controls (**Fig. 1f**).

Collectively, these results demonstrated an oncogenic role of *SMARCB1* loss in EpS, which appears to be conveyed by pleiotropic pathways and reflected in widespread changes to the cellular transcriptome.

### Residual SWI/SNF complex conserves chromatin accessibility in a subgroup of physiological SWI/SNF sites

Since genes regulating epigenetic modifications showed altered expression upon *SMARCB1* re-expression (**Fig. 1c**) and *SMARCB1* is known to participate in SWI/SNF mediated enhancer regulation^25^ we next investigated the functionality of the (residual) SWI/SNF with regard to chromatin remodeling^25,28,29^. Reconstitution of the physiological SWI/SNF complex following re-expression of *SMARCB1* may thus act by modulating SWI/SNF chromatin remodeling, thereby disrupting tumor-maintaining properties of the residual complex. To explore the function of the residual SWI/SNF complex on chromatin remodeling in EpS, we first carried out genome-wide footprinting for open chromatin by Assaying for Transposase-Accessible Chromatin using Sequencing (ATAC-Seq) in four cell line models from **Fig. 1a** (FU-EPS-1, HS-ES-2M, NEPS, VA-ES-BJ) representing both subtypes (proximal and distal). As depicted in **Fig. 2a**, re-expression of *SMARCB1* had profound effects on chromatin accessibility in all EpS cell lines tested, indicating that *SMARCB1* re-expression may indeed affect the affinity and function of the residual SWI/SNF complex. DiffBind analyses across all EpS cell lines further suggest that *SMARCB1* re-expression considerably increased chromatin accessibility while SWI/SNF inhibition via treatment with BRM014 (also known as Compound 14), a small molecule dual inhibitor of both SWI/SNF-ATPases BRG1 and BRM^26,33^, resulted in mostly lost chromatin accessibility (**Supp. Fig. 2a**). To further elaborate on the mechanisms of the observed functional shift, we conducted chromatin immunoprecipitation followed by DNA-sequencing (ChIP-Seq) experiments in VA-ES-BJ as a representative cell line against the ATPase subunit BRG1 as well as SMARCB1 to directly map the genomic interactions of the (residual) SWI/SNF complex. In parallel, we probed for histone marks indicative for active enhancers (H3K27ac), active promoters (H3K4me3), and polycomb repression (H3K27me3) to elucidate the functional chromatin status after reconstitution of the physiological SWI/SNF complex. Here, we found that *SMARCB1* re-expression led to prominent changes in SWI/SNF binding as well as active histone marking (H3K27ac and H3K4me3) in accordance with gene signatures detected by Affymetrix (**Fig. 2b**). As shown in **Fig. 2c**, differentially accessible chromatin regions showed gains of H3K27ac occupancy upon re-expression of *SMARCB1*, thus likely constituting SWI/SNF specific *cis*-regulatory elements (sCRE) that drive the observed functional characteristics and changes in EpS after regaining active enhancer histone marks. Likewise, BRG1 occupancy (as a proxy for (residual) SWI/SNF) that appears to be restricted to an enriched subgroup of physiological SWI/SNF sites in the wild-type condition readily regains a broader distribution across sCRE sites after re-expression of *SMARCB1*. The overall low enrichment of BRG1 at and around regulated gene transcription start site (TSS) indicated that a majority of (residual) SWI/SNF function may be tied to modulation of chromatin accessibility at distal sCREs (mostly 50–500 kb from TSS) rather than directly opening chromatin at proximal sCREs (**Supp. Fig. 2a**). Simple motif Enrichment Analyses (SEA) of differentially accessible chromatin regions showed that enriched motifs found in binding sites lost upon BRM014 treatment and gained upon *SMARCB1* re-expression appear to be from similar groups of transcription factor families (**Fig. 2d**). Sites gained after BRM014 treatment or lost upon DOX treatment were too few to conduct motif analysis (**Supp. Fig. 2a**). The enrichment ratio of AP-1 family transcription factors (e.g. JUND, FOSL1) was greatly increased after *SMARCB1* re-expression, while differentiation and development associated transcription factors (e.g. VEZF1, KLF8) showed less striking but nevertheless notable increases in enrichment ratio. These findings are mirrored in SEA of differentially bound BRG1 sites with/without SMARCB1 re-expression (**Fig. 2d**). Overall, this is in agreement with prior data published on the interactome of SWI/SNF demonstrating the essential role of AP-1 family transcription factors as pioneer factors facilitating epigenetic restructuring by cooperation with the SWI/SNF complex^21,34^. Differential SEA comparing motifs enriched in the chromatin regions lost upon BRM014 treatment with regions gained upon DOX treatment show motif enrichment of transcription factors regulating cell cycle progress and apoptosis downregulated upon BRM014 treatment, while *SMARCB1* re-expression was associated with enrichment of developmental and homeobox transcription factors (**Supp. Fig. 2b**).

**Fig. 2:**
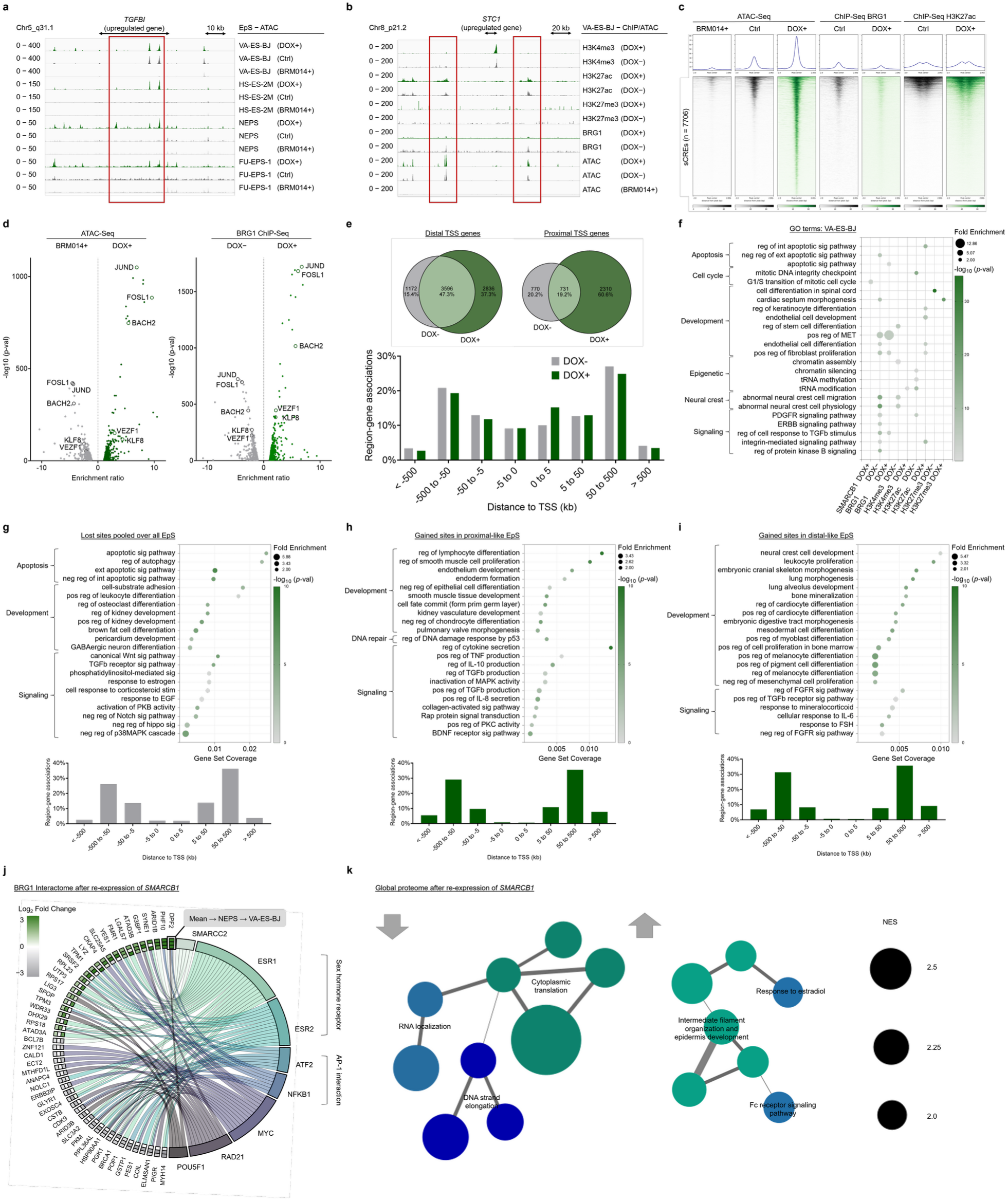
(a) Example ATAC-Seq tracks of EpS cell lines at the TGFBI site showing opened chromatin in the DOX+ condition and closed chromatin in the BRM014+ condition. (b) Example ChIP-Seq and ATAC-Seq tracks in VA-ES-BJ at the STC1 gene site demonstrating differential histone marking and open chromatin distribution correlating with upregulated gene expression in transcriptomic data. (c) Heatmaps of BRG1 and H3K27ac (active enhancer mark) occupancy as well as chromatin accessibility across loci differentially enriched in ATAC-Seq. (d) Volcano plots of Simple Enrichment Analysis (SEA) demonstrating highly enriched motifs in open chromatin regions lost upon BRM014 treatment and gained upon SMARCB1 re-expression (pooled across all EpS cell lines) as well as at BRG1 ChIP sites with and without BAF47 re-expression (in VA-ES-BJ). (e) GREAT region-gene-association distribution of BRG1 sites binned by distance to closest gene TSS and Venn diagrams of associated genes proximal (≤ 2kb) and distal (up to 1.000 kb) from TSS. (f) DiffBind GREAT analysis of differentially regulated BAF47 and BRG1 sites as well as histone marks upon re-expression of SMARCB1, similar GO terms were summarized. (g) DiffBind GREAT analysis of sites of lost chromatin accessibility upon treatment with BRM014 alongside GREAT region-gene-associations binned by distance to closest gene TSS. (h-i) DiffBind GREAT analysis of sites of gained chromatin accessibility upon re-expression of SMARCB1 in proximal- and distal-like EpS cell lines alongside GREAT region-gene-associations binned by distance to closest gene TSS. (j) Chord diagram showing the gene names of proteins co-immunoprecipitating with the core SWI/SNF-ATPase BRG1 significantly regulated in NEPS and VA-ES-BJ upon SMARCB1 re-expression as different annuli of log2 fold changes on the left (from outermost to innermost: Mean, NEPS, VA-ES-BJ) with Enrichr-based curated transcription factor protein-protein interactions shown on the right. Left–right connections indicate gene/protein membership in the transcription factor’s interactome. (k) GSEA-based network analysis of up-/down-regulated biological process sets upon SMARCB1 re-expression from shared regulated proteins in VA-ES-BJ and NEPS.

Genomic regions enrichment of annotations tool (GREAT) analyses suggested that the number of BRG1 associated genes was greatly elevated after re-expression of *SMARCB1*. The distribution of gained BRG1 ChIP-Seq loci included a higher percentage of close proximity (0–5 kb) sites downstream of the TSS of putatively regulated genes with a greater percentage of gained proximal (≤ 2kb from TSS) than distal (up to 1 Mb from TSS) gene associations, possibly due to restoration of *SMARCB1*-associated mitotic bookmarking function promoting affinity to TSS regions^16^ (**Fig. 2e**). On the other hand, sCRE occurred almost entirely at distal enhancer regions ( **Fig. 2c, Supp. Fig. 2a**). Furthermore, compared with the proliferation associated motifs enriched in residual SWI/SNF binding loci (BRG1 DOX−), motifs of SMARCB1 sites (SMARCB1 DOX+) appeared to be more development-associated (**Supp. Fig. 2c**). This epigenetic shift may in part represent the re-activation of lineage-dependent development pathways via the bookmarking function of SMARCB1 previously described^16^. Overall, these findings are in line with previous studies of the SWI/SNF complex postulating the necessity of constantly active SWI/SNF-driven chromatin modulation to retain accessibility to enhancer regions which in turn may facilitate transcription factor binding associated with tumorigenic expressional programs^27,35,36^. Moreover, consistent with previous studies characterizing the promoter categories of the unchanged/retained compared with the gained BRG1 associated genes upon re-expression of *SMARCB1*^23^, we also found an increased percentage of bivalent and polycomb-repressed (H3K27me3) promotors at gained distal (up to 1 Mb from peak) but not proximal (≤ 2 kb) BRG1 associated genes (**Supp. Fig. 3a**). Alongside SWI/SNF modulation shown at sCREs, this may be indicative of epigenetic restructuring occurring predominantly at distal sites. GREAT gene ontology (GO) term analysis of differentially bound histone marks and BRG1 loci showed that associated genes are most strongly implicated in the regulation of apoptosis, cell cycle progress, development as well as several proliferation and differentiation associated hormonal signaling pathways (**Fig. 2f**). While global chromatin accessibility correlated strongest within the respective EpS cell line models (with similar subtypes clustering together more closely), chromatin accessibility patterns at sCRE sites were most similar within the BRM014 treatment condition, regardless of cell line (**Supp. Fig. 3b**). On the one hand, this indicates that residual SWI/SNF regulation at a subgroup of sCRE may constitute a specific collection of loci highly conserved across EpS, largely irrespective of histological subtype, possibly intricately involved in tumor maintenance. On the other hand, global chromatin accessibility patterns appear to be more closely tied to individual cell line and histological subtype. This hypothesis further suggests that HS-ES-2 cell lines which lacks histological metadata (HS-ES-2R: unknown relapse, HS-ES-2M: lung metastasis, both from the same patient) are derived from the proximal subtype. This is also in line with morphological and functional characteristics observed in HS-ES-2 which most closely resemble VA-ES-BJ as a confirmed proximal EpS cell line (**Supp. Fig. 3b**).

Based on these clustering results, we performed GREAT-based GO terms analyses of the sCRE sites within similar subgroups. The sCRE sites lost upon BRM014 treatment (pooled across all EpS lines) appeared to be enriched in gene sets most strongly associated with the regulation of apoptosis while gained sites upon re-expression of *SMARCB1* showed subtype-dependent signaling and development-associated gene sets (**Fig. 2g-i**). Only regulation of and response to TGFb pathway signaling seems to be preserved across all sCRE groups, potentially highlighting it as a central pathway in EpS. The abrogation of pro-oncogenic, anti-apoptotic residual SWI/SNF function and retargeting towards cell development and differentiation may thus cooperate to reduce tumor viability.

Next, to confirm regulated pathways predicted by our epigenetic analyses, we performed mass spectrometry-based quantification of nuclear proteins co-immunoprecipitated with SWI/SNF-ATPase BRG1 in two representative cell lines (NEPS and VA-ES-BJ). Enrichr-based curated transcription factor protein-protein interaction gene set enrichment analysis^37,38^ demonstrated that *SMARCB1* re-expression increased interaction with other SWI/SNF subunits (e.g. ARID1B), indicating that SWI/SNF assembly might be influenced by *SMARCB1* status (**Fig. 2j**). The influence of the AP-1 transcription factor family and its association with sex hormonal pathways, as predicted by ATAC-Seq, is also evident in the regulated interactomes of ATF2 and NFKB1 (ref. ^39^), as well as the estrogen receptors ESR1 and ESR2. Other differentially regulated interactome sets upon *SMARCB1* re-expression point towards TFs related to proliferation, chromatin organization and cell fate determination (e.g. MYC, RAD21, POU5F1) (**Fig. 2j**). Furthermore, GSEA analysis shows that several gene sets pertaining to chromosome organization and telomere maintenance are downregulated upon re-expression of *SMARCB1* while upregulated gene sets are mostly centered around energy metabolism and development (**Supp. Fig. 4**).

GSEA based WGCNA analysis of whole cell global proteomics confirm prior predictions based on transcriptomic and epigenetic data, wherein we show the downregulation of proteins associated with DNA/RNA regulation and translation while development and cytoskeleton associated proteins are upregulated upon restoration of the physiological SWI/SNF (**Fig. 2k**). Interestingly, increased response to estradiol was also found to be enriched, mirroring the increased interaction with the sex hormone signaling pathway seen in interactome analyses (**Fig. 2l, 2k**).

### Residual SWI/SNF complex as a druggable target for EpS

Since these results provided further evidence that the residual SWI/SNF complex may constitute a promising druggable target, we performed clonogenic growth assays and drug-response assays (**Fig. 3a. 3b, Supp. Fig. 5a-i**) with BRM014. As shown in **Figs. 3a** and **3b**, treatment of EpS cell lines with BRM014 reduced proliferation and clonogenic growth in a dose-dependent manner. Interestingly, these effects were rather weak in short-term assays (48h Resazurin metabolization assays) (**Supp. Fig. 5j**) but became more prominent in long-term assays (2-week clonogenic assays) (**Fig. 3a, 3b, Supp. Fig. 5a-i**), which is consistent with a delayed effect based on epigenetic remodeling. As it has been demonstrated before^27^, the cytotoxic effects of SWI/SNF ATPase inhibition appears particularly enhanced in SWI/SNF-dependent/deficient entities while displaying good *in vivo* tolerability at effective doses^33^ (**Fig. 3a**). This trend towards targeted action is mirrored in our investigations with all EpS cell lines exhibiting IC_50_ values well below the reported AAC_50_ value of 10 nM^33^ while *SMARCB1*-proficient control cell line IC_50_ values were all above (**Fig. 3b**).

**Fig. 3:**
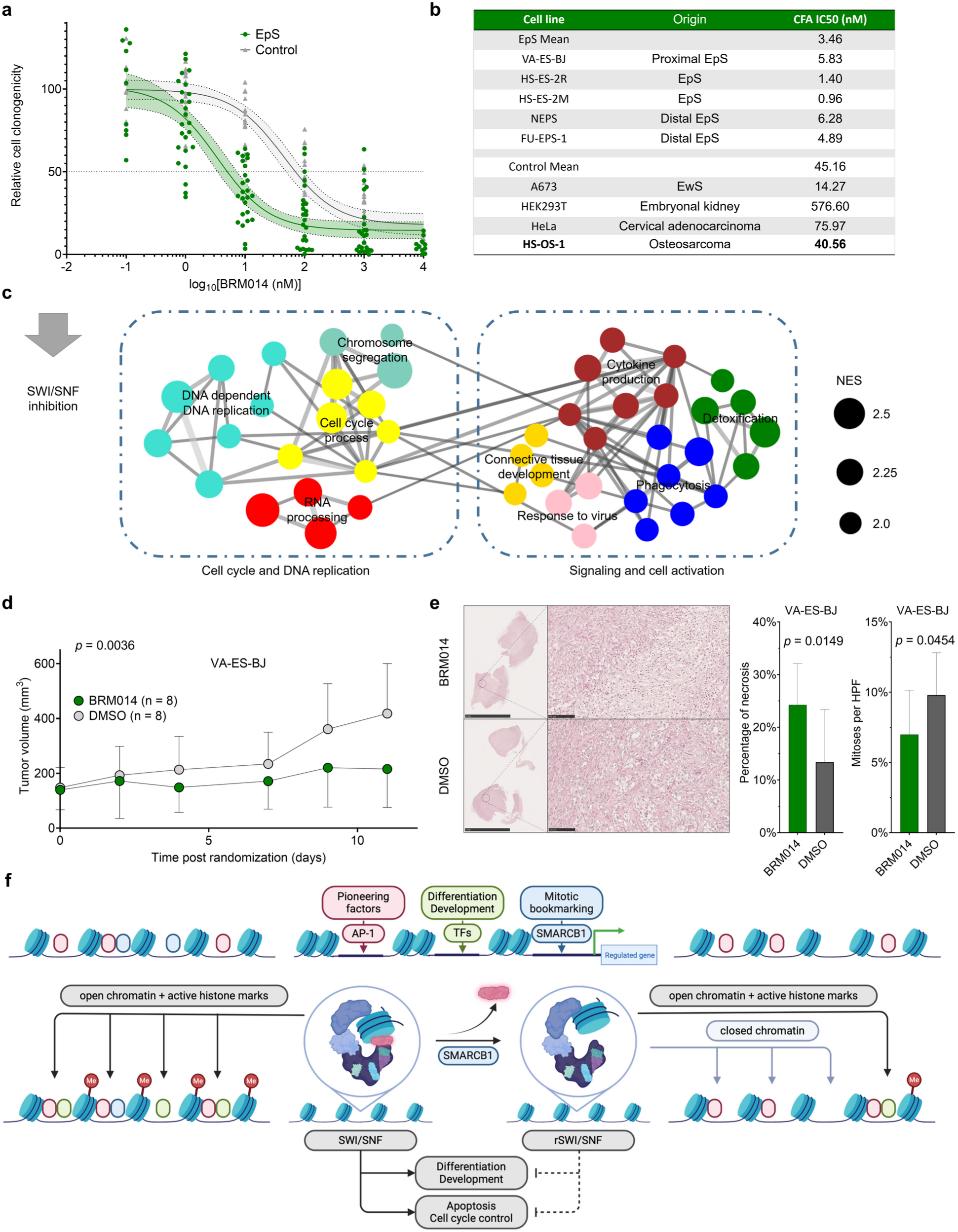
(a) Colony formation assay and Brightfield images of VA-ES-BJ demonstrating synergistic action of SMARCB1 re-expression and SWI/SNF ATPase inhibition (via BRM014). (b) Pooled BRM014 drug response curves with 95% CI and individual calculated IC50 values of SWI/SNF-deficient EpS cell lines (FU-EPS-1, HS-ES-2M, HS-ES-2R, NEPS, VAESBJ) plotted against SWI/SNF wild-type control cell lines (HEK293T, HeLa, A673, HS-OS-1) based on clonogenicity assay. (c) GSEA-based network analysis of down-regulated gene sets upon rSWI/SNF inhibition by BRM014 treatment from shared regulated genes in HS-ES-2M, HS-ES-2R and NEPS. (d) Schematic for BRM014 *in vivo* experiment alongside growth curves of VA-ES-BJ xenografts under BRM014 treatment or vehicle control (DMSO). (e) Example of cross-sectional VA-ES-BJ xenograft H/E stains after treatment with BRM014 or DMSO vehicle control and bar plots of percentage of necrosis and number of mitoses per HPF observed in xenograft histology in the BRM014 vs. DMSO condition in VA-ES-BJ. (f) Schematic of (r)SWI/SNF action in EpS.

As a control for assembly-specific SWI/SNF interactions, we performed drug assays with the PROTAC dBRD9 (ref. ^40^). dBRD9 selectively degrades BRD9 – a core subunit exclusive to the ncBAF complex which is the only main SWI/SNF assembly known not to contain SMARCB1 and which has been demonstrated to be vital for tumor maintenance in other SWI/SNF-mutant entities^13,41–44^. Contrary to these entities (MRT, synovial sarcoma, multiple myeloma), our drug-response assays using dBRD9 alongside BRM014 CFAs in Eps cell lines demonstrated that the SMARCB1 incorporating cBAF and PBAF complexes with intact ATPase function but not the SMARCB1 independent ncBAF complex appeared to be crucial to tumor survival in EpS (**Supp. Fig. 5k**).

In line with our epigenetic and functional results, GSEA-based WGCNA analysis of our transcriptomic data prominently showed enriched downregulated gene sets mostly associated with cell cycle and DNA replication as well as cell activation and signaling (**Fig. 3c**), while only one upregulated gene set (negative regulation of muscle cell differentiation, NES = 2) was identified.

To evaluate the *in vivo* therapeutic viability of SWI/SNF ATPase inhibition as a novel targeted approach for EpS, we conducted drug trials in EpS s.c. xenograft NSG mice models by intraperitoneal (i.p.) injection of BRM014 (20 mg/kg, 5 days/week for 2 weeks^45,46^). BRM014 treatment significantly suppressed tumor growth with minimal observed general toxicity (**Fig. 3d**). Histology of the BRM014 treated xenografts show a significantly decreased mitosis index alongside increased percentage of observed necrosis when compared to DMSO vehicle controls (**Fig. 3e**). These results are concurrent with mechanistic insights demonstrated in our epigenetic and transcriptomic as well as previously published data^16,21^ (**Fig. 3f**). Contrary to initial expectations, our findings highlight that targeting SWI/SNF represents a promising therapeutic approach for treating EpS despite its hallmark SWI/SNF deficiency. This discovery may prompt further investigations into whether similar targeted strategies could be effective in the multitude of SWI/SNF deficient entities.

## DISCUSSION

To date, exact mechanisms driving EpS have remained elusive, with some previous studies proposing a loss of function of the SWI/SNF complex upon deletion of *SMARCB1* and the subsequent destabilization of the epigenome as the main driver of EpS^10^. However, our findings − while fully supporting the core role of *SMARCB1* in EpS − also bring forward the possibly counterintuitive notion that loss of *SMARCB1* may convert the canonically tumor-suppressive SWI/SNF complex into an oncogenic residual complex. In fact, we show that this aberrant residual complex and its immediate interaction partners assist in maintaining EpS in a state of anti-apoptotic and proliferative de-differentiation, hence constituting an ideal target for mechanistic therapy. Targeting of the SWI/SNF ATPase, already proven to be effective and of low off-target toxicity^27,33^ in other tumor entities, is considered as promising novel targeted therapy^26^. Accordingly, approaches involving radiation^24^ and epigenetic modulation have already been successful for the treatment of EpS, possibly due to interaction with key downstream patho-mechanistic players of *SMARCB1*-deficient SWI/SNF, such as HDAC^47,48^. As BRM014 treatment seems to disrupt rSWI/SNF mediated tumor maintenance, combination therapies including BRM014 may likely work synergistically for EpS treatment.

Importantly, we demonstrate the complexity of perturbations affecting the SWI/SNF complex which exhibits canonical traits of both traditional tumor suppressors and proto-oncogenes and may be translatable to the multitude of other SWI/SNF-dependent entities. Mechanistically, the residual SWI/SNF appears to retain partial function of the physiological SWI/SNF at a conserved group of *cis*-regulated elements, primarily exhibiting pro-proliferative and anti-apoptotic activity while lacking sufficient differentiation and development promoting features. We show that this vulnerability can be exploited by either abrogating residual SWI/SNF activity entirely or by rescuing physiological SWI/SNF functionality through SMARCB1 re-incorporation. The latter mechanism has also been shown to be partially viable through targeting physiological degraders (residual) SWI/SNF in other SWI/SNF-mutant tumor entities^34^. Recently, an orally available selective inhibitor of BRG1/BRM (FHD-286) has shown both preclinical and clinical therapeutic promise against hematological entities^49,50^.

Interestingly, both, the restoration of physiological SWI/SNF as well as the inhibition of residual SWI/SNF led to a downregulation of immature cell types in the transcriptome (**Supp. Fig. 6a, 6b**) which mirrors the major regulated pathways derived from our ATAC-Seq data. Furthermore, abnormal, neural crest cell lineage associated GO terms highly enriched in ChIP-Seq are down-regulated upon initiation of fibroblast/keratinocyte-associated developmental pathways concurrent with the re-expression of *SMARCB1* (**Fig. 2f**). This may suggest the maintenance of an undifferentiated neural-crest-like stem cell state in wild-type EpS, both as a mechanism for oncogenesis as well as a possible phylogeny. A similar origin has already been suggested for MRT^17^. Altered TGFb pathway signaling which has been tied to cellular differentiation, growth control and EMT processes^51,52^ observed in our transcriptomic and epigenomic data could be mechanistically linked to the morphological and functional changes in EpS cell lines upon SMARCB1 re-expression and SWI/SNF inhibition (**Supp. Fig. 6c**). Both show a synergistic effect decreasing clonogenicity as they act through slightly different mechanisms. When both treatments are administered, some rSWI/SNF are inhibited by BRM014 while other rSWI/SNF may instead re-incorporate SMARCB1 and regain physiological function. Given that TGFb signaling exhibits a context-dependent role, acting as an early suppressor and a late promoter of tumorigenesis ^52^, this dual functionality may similarly underpin the mechanistic versatility of the SWI/SNF complex as suggested by our data.

Collectively, our results identify the dependency on the residual SWI/SNF as an intrinsic vulnerability for targeted therapy in *SMARCB1*-deficient entities. This may provide a starting point for further research to uncover whether this abnormal, tumor-maintaining SWI/SNF state is shared amongst the spectrum of SWI/SNF-dysfunctional entities but possibly triggered by different mechanisms.

## METHODS

### Provenience of cell lines and culture conditions

Human Epithelioid Sarcoma (EpS) cell lines were obtained from the following repositories/providers: HE-ES-1, HS-ES-2M, and HS-ES-2R from RIKEN cell bank (Japan); VA-ES-BJ from the German Collection of Microorganisms and Cell cultures (DSMZ, Braunschweig, Germany). NEPS from Niigata University (Japan); FU-EPS-1 (ST257) from Fukuoka University (Japan); and Epi-544 from MD Anderson Cancer Center, University of Texas (Houston, USA). All EpS cell lines exhibited virtually no detectable *SMARCB1* expression (Ct_q_ values > 35 in TaqMan qPCR). Human Ewing sarcoma (EwS) cell lines were obtained from the following repositories/providers: A-673 from American Type Culture Collection (ATCC); MHH-ES1 and SK-N-MC from DSMZ, TC-106 and TC-71 from the Children’s Oncology Group (COG). Human osteosarcoma cell line HS-OS-1 was obtained from RIKEN cell bank (Japan). Human HEK293T and HeLa cells were obtained from DSMZ. All cell lines were cultured in RPMI 1640 with stable glutamine (Gibco) supplemented with 10% fetal bovine serum (FBS), 100 U/ml penicillin and 100 μg/ml streptomycin (Sigma-Aldrich) at 37°C in a fully humidified 5% CO2 atmosphere. Cell lines were routinely tested for Mycoplasma contamination using a custom nested PCR protocol. Cell line identity was confirmed by STR and/or SNP profiling.

### Extraction of total DNA and RNA, reverse transcription, and quantitative Real-Time PCR (qRT-PCR)

Total DNA extraction was performed using NucleoSpin Tissue mini kit (Macherey-Nagel, Germany). RNA was extracted using NucleoSpin RNA mini kit (Macherey-Nagel) including a 15 min DNAse-treatment, and reversely transcribed using the High-Capacity cDNA Reverse Transcription Kit (Applied Biosystems). qRT-PCRs were performed in a final volume of 15 µl using SYBR Select Master Mix (Applied Biosystems) or TaqMan Fast Advanced Mastermix (Thermo Fisher). All primer sequences used for qRT-PCR are listed in **Supplementary Table 1**. Cycling conditions were as follows: 50°C for 2 min (UNG/UDG incubation), 95°C for 2 min (polymerase activation), then 40 cycles at 95°C for 15 sec (denaturation) and 60°C for 1 min (annealing, elongation, and detection).

### Cloning, plasmid design, and lentiviral transduction

Cloning and plasmid design was performed in cooperation with the Cellular Tools Core Facility at the DKFZ, using a pENTR223-*SMARCB1* entry vector to recombine the *SMARCB1* coding sequence into the doxycyline inducible expression vector rwpSMART-Tre3G-GW-mCMV-Puro-2A-TetON3G (CellularTools CF) expression plasmid backbone via Gateway technology (ThermoFisher, Braunschweig, Germany). Correct insertion into the plasmids was verified by Sanger-sequencing and agarose gel electrophoresis. Verified, endotoxin-free plasmids were then used for lentiviral transduction: Briefly, HEK293FT cells (ThermoFisher) were co-transfected with the lentiviral constructs (rwpSMART-Tre3G-GW-mCMV-Puro-2A-TetON3G with the *SMARCB1* ORF containing a stop codon) and second-generation viral packaging plasmids VSV.G (Addgene #14888) and psPAX2 (Addgene #12260). 48 h after transfection, the supernatant containing the virus particles was collected, cleared by centrifugation (5 min/500 g) and passaged through a 0.45 μm filter to remove remaining cellular debris. EpS cells were transduced with lentiviral particles at 80% confluency in the presence of 10 μg/mL polybrene (Merck, Germany) for 24 h. For viral transduction only Eps cell lines without detectable SMARCB1 protein expression and characteristics of homozygous SMARCB1-deletion (FU-EPS-1; HS-ES-1, -2M, -2R; NEPS; VA-ES-BJ) were selected. Transduced cells were selected in the 1 μg/ml puromycin (Invivogen) for ∼1 week. Stably transduced isogenic cell lines were derived from single-colonies and characterized for mRNA expression by TaqMan qRT-PCR.

Dox levels for *SMARCB1* re-expression cell lines were titrated to quasi-physiological levels. TaqMan qPCR was performed with EwS cell lines (A-673, MHH-ES1, SK-N-MC, TC-106, and TC-71) which do not show perturbation of the *SMARCB1* gene, alongside *RPLP0* as a known housekeeping gene in EwS and cross-referenced with nTPM values obtained from the Human Protein Atlas (HPA; www.proteinatlas.org)^53^. Across EwS cell lines, a rather narrow variance of *SMARCB1* expressional status could be observed that represented the lower end of the range of *SMARCB1* expression found in non-cancerous cell lines obtained from the HPA (https://www.proteinatlas.org/ENSG00000099956-SMARCB1/cell+line#non-cancerous). As the cell of origin is unknown for EpS, *SMARCB1* expression was adjusted to these levels instead.

### Western blot

Western blots were performed as previously described^54^. For preparation of protein lysates, 4–5×10^5^ cells depending on the cell line were seeded per well into 6-well plates to reach 80% confluency after 72 h (∼1×10^6^ cells). Thereafter, medium was removed, cells were washed with 1 ml of PBS and lysed by adding 100 µl of RIPA buffer (Serva electrophoresis, Heidelberg, Germany) supplemented with cOmplete, Mini, EDTA-free Protease Inhibitor Cocktail (Roche) and PhosStop (Roche). Antibodies were used to detect SMARCB1 (Cell Signaling, #91735, 1:1,000) or GRP94 (Cell Signaling, #20292, 1:1,000) by reaction with a secondary HRP-conjugated, monoclonal murine, anti-rabbit antibody (sc-2357 1:2,000, Santa Cruz).

### Clonogenic growth assays

For clonogenic growth assays, *SMARCB1* re-expressing EpS cells and respective controls were seeded in triplicate wells at low density (5×10^2^ cells) per well in 12-well plates and grown for 10– 14 d (depending on the cell line) with/without DOX-treatment (renewal of DOX or vehicle every 48 h). Thereafter, colonies were stained with crystal violet (Sigma-Aldrich) for visualization of clonogenicity using ImageJ.

### Cell cycle analysis by flow cytometry

Prior to analysis, cells were harvested after 96 h treatment with DOX (refreshed every 48 h) by centrifugation at 1,200 rpm for 4 min, then washed with PBS and spun down at 2,000 rpm for 4 min to collect cells. Excess medium was carefully aspirated afterwards without disturbing the cell pellet, leaving about 100 µl in flask. To fix the cells, the pellet was vortexed while adding 1 ml of ice-cold EtOH (70%) dropwise and stored at 4 °C until analysis. For flow cytometric cell cycle analysis via propidium iodide (PI)-staining, fixed samples were centrifuged at 2,000 rpm for 10 min, then washed twice with cold PBS. Excess medium was aspirated after every washing step. Each sample was resuspended in 500 µl of PI staining buffer (470 µl FACS staining buffer (BioLegend, CA, USA) + 25 µl PI solution (BioLegend) + 5 µl RNase A (VWR International, Pennsylvania, USA) and incubated in the dark at room temperature. Analysis with BD FACS Canto II system was performed within 3 h after staining on slow/medium flow mode. *.fcs output files were gated by FSC-A; SSC-A to define population by size and granulation and by FSC-A; FSC-H/W to rule out doublets. Single cell populations were then analyzed by PI stain in FlowJo (v10) to determine the counts for each cell cycle phase (G1/G0, S, G2/M).

### Transcriptome analyses

To assess the effect of *SMARCB1* rescue on gene expression in EpS cells, microarray analyses were performed. For this, 5×10^4^ cells per well were seeded in 6-well plates and treated with 0.1 μg/μl DOX for 96 h (DOX-refreshment after 48 h). Thereafter, total RNA was extracted with the NucleoSpin RNA mini kit (Macherey-Nagel) and RNA quality was assessed with a TapeStation 4150 system on an RNA Screentape. All samples had an RNA integrity number (RINe)>9 and were hybridized to Human Affymetrix Clariom D microarrays. Gene expression data were quantile normalized with Transcriptome Analysis Console (v4.0.2; Thermo Fisher Scientific) using the SST-RMA algorithm as previously described^55^. Annotation of the data was performed using the Affymetrix library for Clariom D Array (version 2, *Homo sapiens*) on gene level. Differentially expressed genes (DEGs) with consistent and significant fold changes (FCs) across cell lines were identified as follows: i) Normalized gene expression signals were log2 transformed; ii) to avoid false discovery artifacts due to the detection of only minimally expressed genes, all genes with an equal or lower expression value than that observed for *SMARCB1* of the control cell lines (all with homozygous *SMARCB1* deletion) across all cell lines and conditions were excluded. The FCs of the empty vector samples and all *SMARCB1* re-expressing EpS cell lines after treatment (DOX or BRM014) were calculated for each cell line separately. Then the FCs in the *SMARCB1* re-expressing samples were normalized to that of the empty vector cells (for DOX treatment). Finally, resulting FCs were filtered for genes equally down- or up-regulated across cell lines, then averaged to obtain the mean FC per gene. DEGs were determined as having a |log_2_FC| > 0.5.

### Gene-set enrichment analysis (GSEA) and WGCNA

To identify enriched gene-sets, genes were ranked by their expression FC between treatment and control groups, then additionally filtered for unidirectionally aligned regulation in all cell lines for pooled analyses (i.e. up-/downregulated in all cell lines upon DOX treatment). GSEA (multilevel) was performed using the FGSEA R package (v 3.6.3) based on Gene Ontology (GO) biological processes and cell signature terms from MSigDB (c5.go.bp.v7.5.1.symbols.gmt and c8.all.v2023.2.Hs.symbols.gmt)^56^. GO terms were filtered for statistical significance (adjusted *p-* value < 0.05) and a normalized enrichment score |NES| > 2. To construct a network, the Weighted Gene Correlation Network Analysis R package (WGCNA R)^57^ was used. Briefly, a binary matrix of GO-terms × genes (where 1 indicates the gene is present in the GO term and 0 indicates it is not) was created. Then, the Jaccard’s distance for all possible pairs was computed to create a symmetric GO adjacent matrix. Clusters of similar GO terms were identified using dynamicTreeCut algorithm, and the top 20% highest edges were selected for visualization. The most meaningful among highest scoring node in each cluster was determined as the cluster label. The obtained network and nodes files were processed via Cytoscape (v 3.8.0) for network design and visualization as previously described^58^. GOChord visualization for the most highly enriched gene sets/cell signatures was performed with the GOPlot R package^59^. Only genes contained within leadingEdge analysis of plotted cell signature gene sets were shown.

### Protein sample preparation and immune-precipitation followed by mass spectrometry

Nuclear and whole-cell extracts were prepared from 1.5×10^7^ cells per condition with and without DOX treatment for 4 d using the Nuclear Extract Kit (Active Motif). Co-immunoprecipitation was performed with the nuclear fraction using the Dynabeads Co-Immunoprecipitation Kit (Thermo Fisher) coupled with the anti-BRG1 antibody ab110641 (rabbit monoclonal, Abcam), according to the manufacturer’s manual. Mass spectrometry was performed at the DKFZ Core Facility for Genomics and Proteomics as follows:

Proteins (5 µg) were run for 0.5 cm into an SDS-PAGE and the entire piece was cut out and digested using trypsin according to Shevchenko et al.^60^ adapted to on a DigestPro MSi robotic system (INTAVIS Bioanalytical Instruments AG). A LC-MS/MS analysis was carried out on a Vanquish Neo UPLC (Thermo Fisher) directly connected to an Orbitrap Exploris 480 mass spectrometer for a total of 90 min. Peptides were online desalted on a trapping cartridge (Acclaim PepMap300 C18, 5 µm, 300 Å wide pore; Thermo Fisher) using the metering device at a flow rate of 30 µl/min for total loading volume of 60 µl. The analytical multistep gradient (300 nl/min) was performed using a nanoEase MZ Peptide analytical column (300 Å, 1.7 µm, 75 µm x 200 mm, Waters) using solvent A (0.1% formic acid in water) and solvent B (0.1% formic acid in acetonitrile). For 72 min the concentration of B was linearly ramped from 4% to 30%, followed by a quick ramp to 80%, after four minutes the concentration of B was lowered to 2% and a three column volume equilibration step was appended. Eluting peptides were analyzed in the mass spectrometer using data dependent acquisition (DDA) mode. A full scan at 60k resolution (380– 1,400 m/z, 300% AGC target, 45 ms maxIT) was followed by up to 1.5 s of MS/MS scans. Peptide features were isolated with a window of 1.4 m/z, fragmented using 26% NCE. Fragment spectra were recorded at 15k resolution (100% AGC target, 54 ms maxIT). Dynamic exclusion was set to 30 s. Data analysis was carried out by MaxQuant (version 2.1.4.0, ref. ^61^) using an organism specific database extracted from Uniprot.org (human reference database with one protein sequence per gene, containing 20,597 unique entries from ninth of February 2024). Settings were set to default with the following adaptions. Match between runs (MBR) was enabled to transfer peptide identifications across Raw files based on accurate retention time and m/z. Fractions were set in a way that MBR was only performed within replicates. Separate parameter groups were assigned for cell line and proteome fraction or IP respectively. Separate Label free quantification (LFQ) per parameter group was enabled. Quantification was performed using the LFQ approach based on the MaxLFQ algorithm^62^. A minimum of 2 quantified peptides per protein was required for protein quantification. In addition, iBAQ-values^63^ were generated.

Label-free quantification results were then normalized to column sum per experimental subgroup and missing values imputed using slsa (for partially observed values) and det quantile (quantile 2.5, factor 1, for values missing in the entire collection) before hypothesis testing in Prostar^64^. Resulting FCs were filtered for genes/proteins equally down- or up-regulated across cell lines, then averaged to obtain the mean FC per gene. GSEA enrichment analysis followed by WGCNA was then performed on the prefiltered gene/protein lists as detailed in the GSEA/WGCNA section. Gene sets shown in visualizations were filtered for |NES| > 2.5 (downregulated upon *SMARCB1* re-expression in BRG1 coIP) or |NES| > 2 (all other conditions) to maintain legibility.

Differentially enriched proteins (DEP) were defined with a |log_2_FC| > 1 and adjusted *p*-value < 0.05. Gene set enrichment analyses were performed on DEP from the BRG1 coIP (+/-re-expression of *SMARCB1*) condition using the “Protein-protein interaction (PPI) hub protein” sets within the Enrichr tool^37^. GOChord visualization was performed with the GOPlot R package^59^.

### Assay for transposase-accessible chromatin followed by next-generation sequencing (ATAC-Seq)

Nuclei isolation and sample preparation was performed using the ATAC-Seq kit and the 24 UDI for Tagmented libraries - Set I (Diagenode) with 5×10^4^ cells per condition after treatment with DOX or BRM014 (1 µM) for 4 days, according to the manufacturer’s protocol. In brief, cells were treated, harvested, and then lysed. Nuclei were then isolated and DNA tagmented. Library amplification was performed with unique dual index primers. Size selection was performed on the libraries with AMPure XP beads (Beckman). Finally, quality control was performed with the Qubit DNA Assay and the Agilent TapeStation 4150 system on a D5000 Screentape.

An equimolar multiplex was created from purified libraries, adjusted to 10 nM. Sequencing was performed on a NovaSeq 6000 Paired-End 100bp S4 chip by the DKFZ NSG Core facility. Using the DKFZ galaxy platform, the sequencing reads were pre-processed with TrimGalore! to filter out adapter sequences, then mapped with bowtie2 against the hg19 reference genome and post-processed using Filter BAM to filter out improperly paired reads, mapping quality less than 30 and/or mapping to the mitochondrial chromosome. Duplicate reads were removed using MarkDuplicates. Processed reads were visualized in IGV using bamCoverage. Peaks were called using MACS2 on the BAM datasets. Differential peak enrichment analysis between treatment conditions and control was performed with the DiffBind R package^65^. Motif analysis on differential peaks was performed with the MEME suite (MEME-ChIP and SEA) on genomic sequences extracted from peak files. Differential motif analysis was performed with SEA using the respective other condition as background sequences (BRM014 associated sequences as background for DOX/*SMARCB1* re-expression associated sequences and vice versa). Discovered differential peaks were filtered by fold enrichment (>2 for BRM014 treatment pooled analysis, >3 for DOX treatment subtype analysis), then analyzed with the GREAT (Ver 4.0.4) tool developed by Stanford University^66,67^. Every gene was assigned a basal regulatory domain 5 kb up- and 1 kb downstream of its TTS, extended distally by a maximum of 1 Mb into each direction without overlapping with the basal domain of the next closest gene, then regions were associated with all their immediately overlapping genes.

Correlation matrices were computed with the ComputeMatrix tool using the whole genome summarized into 10 kb bins or DiffBind site bins respectively. The matrices were then analyzed with Graphpad Prism for Pearson correlation.

### Chromatin immunoprecipitation followed by DNA sequencing (ChIP-Seq)

H3K4me3, H3K27ac, H3K27me3 chromatin immunoprecipitation (ChIP) was performed using the iDeal ChIP-Seq kit for histones or iDeal ChIP-Seq kit for transcription factors (Diagenode) using the following ChIP-Seq grade antibodies: C15410003 (rabbit polyclonal, Diagenode) for H3K4me3, ab4729 (rabbit polyclonal, Abcam) for H3K27ac, C15410069 (rabbit polyclonal, Diagenode) for H3K27me3 as well as #91735 (rabbit monoclonal, Cell Signaling) for BAF47 and ab110641 (rabbit monoclonal, Abcam) for BRG1 respectively. 2×10^7^ cells were cross-linked with 1% formaldehyde for 10 min followed by quenching with 125 mM glycine (final concentration) for 10 min at room temperature. Chromatin was isolated by the addition of lysis buffer, and lysates were sonicated to obtain sheared chromatin to an average length of ∼300 bp. ChIP was performed with chromatin of 1 million cells for histone marks and 4 million cells for transcription factors (TFs). The equivalent of 1% of chromatin used for TFs was kept to quantify the input. ChIP was performed overnight at 4 °C on a rotating wheel with 1.4 μg of antibody for H3K4me3, 1 μg for H3K27ac, 2.9 μg for H3K27me3, 1.4 μg for SMARCB1 and 7 μg for BRG1. After ChIP, chromatin was eluted for 30 min on the DiaMag rotator at room temperature in 100 µl iE1 buffer and reverse cross-linked for 4 hours at 65°C with shaking in iE2 buffer. DNA was precipitated and purified with magnetic beads with the IPure v2 kit (Diagenode). Before sequencing, ChIP efficiency was validated by qPCR for each antibody on specific genomic regions using powerSYBR Green Master Mix (Applied Biosystems) and compared for each primer pair to the input DNA. Primers are listed in **Supplementary Table 1.**

For library preparation from ChIP-fragments the MicroPlex Library Preparation Kit v3 was used along with ‘24 UDI for MicroPlex v3 - Set I’ (Diagenode). 10 µl of every sample containing 50 pg of precipitated DNA were used for library preparation according to the manufacturer’s protocol for Template Preparation and Library Amplification. Amplified intermediate libraries were quantified using the Agilent TapeStation 4150 system on a D5000 HS Screentape to check for fragment distribution, unincorporated adapters as well as DNA concentration and to evaluate the necessity for re-amplification in case of inadequate yields. Where necessary, re-amplification was performed for 2–3 cycles following the same settings as the library amplification protocol.

Finished libraries were purified using AMPure® XP beads in a 1:0.9 volume ratio of sample to beads and eluted in 15 µl low TE-buffer. Libraries were then quantified using the TapeStation 4150 system on a D5000 HS Screentape.

An equimolar multiplex was created from purified libraries, adjusted to 10 nM. Sequencing was performed on a NovaSeq 6000 Paired-End 50bp SP chip by the DKFZ NSG Core facility. Using the DKFZ galaxy platform, the sequencing reads were mapped with BWA for medium and long reads (>150 bp) against the hg19 reference genome and post-processed using NGS: SAMtools, Filter SAM or BAM for any read with mapping quality less than 20. Correlation among different targets was assessed using NGS: DeepTools, multiBamSummary. Processed reads were visualized in IGV using bamCoverage. Peaks were called using MACS2 on the BAM datasets. Motif analysis was performed with the MEME suite (MEME-ChIP and SEA) on genomic sequences extracted from peak files. Differential peak enrichment analysis was performed with the DiffBind R package^65^. Resulting BED files were analyzed with the GREAT (Ver 4.0.4) tool developed by Stanford University^66,67^. Every gene was assigned a basal regulatory domain 5 kb up- and downstream of its TTS, extended distally by a maximum of 1 Mb into each direction without overlapping with the basal domain of the next closest gene, then regions were associated with all their immediately overlapping genes. For analysis of promoter categories, H3K4me3 and H3K27me3 histone mark peaks were associated with gene TSS within 2 kb up- or downstream of peaks using GREAT and overlapped with BRG1 peak associated genes.

### Mouse xenograft experiments

For subcutaneous xenograft experiments, 2.5×10^6^ wild-type or pre-transduced VA-ES-BJ or NEPS EpS cells suspended in a 1:1 mix of PBS and Geltrex (LDEV-Free, hESC-Qualified, Reduced Growth Factor Basement Membrane Matrix-5 mL. A1413302, Gibco/LifeTechnologies) were subcutaneously injected into the flank of NSG mice. Tumor growth was measured three times a week using a caliper. Tumor volumes were calculated using the following formula: V = L × W^2^ / 2, where V is the tumor volume, L the largest diameter and W the smallest diameter. When the tumors reached an average volume of 60 mm^3^, mice were randomized in two groups of which one was henceforth treated with 2 mg/ml DOX (Beladox, Bela-pharm, Germany) dissolved in drinking water containing 5% sucrose (Sigma-Aldrich) to induce an *in vivo* re-expression (DOX (+)), whereas the other group only received 5% sucrose (control, DOX (−)). Before tumors of control groups exceeded a maximum length of 15 mm in any dimension or an average volume of 1,500 mm^3^, all mice of the experiment were sacrificed by cervical dislocation. Other humane endpoints were determined as follows: Ulcerated tumors, loss of 20% body weight, constant curved or crouched body posture, bloody diarrhea or rectal prolapse, abnormal breathing, severe dehydration, visible abdominal distention, obese Body Condition Scores (BCS), motoric irregularities, aggressiveness as a sign of pain, automutilation, apathy, self-isolation, maximum observation period of 12 months and/or invasive tumor growth with fluid leakage (e.g. blood, serous fluid), functional impairment, disability, or pain. Animal experiments were conducted under allowance by the government of North Baden (NTP-ID: 00029631-1-6) and in accordance with the 3R principle of animal experiments (replacement, reduction, and refinement), ARRIVE guidelines, recommendations of the European Community (86/609/EEC), and UKCCCR (guidelines for the welfare and use of animals in cancer research).

For the treatment of pre-transduced xenografts, BRM014 (solubilized in 5% DMSO + 10% Kolliphor + 85% (10% beta-CD in sterile water)) was administered via i.p. injection 20 mg/kg, 5 days a week, for a maximum of 2 weeks^45,46^. After extraction of the tumors, a small fraction of each tumor was snap frozen in liquid nitrogen for preservation, while the remaining tumor tissue was fixed in 4%-formalin and embedded in paraffin for immunohistology. Statistical significance of longitudinal growth was calculated with the TumGrowth online tool by Enot et al.^68^.

### Histology

For IHC, 4-μm sections were cut and stained with eosin/hematoxylin (#T.865.3, Mayer, Roth). Slides were scanned on a Nanozoomer-SQ Digital Slide Scanner (Hamamatsu Photonics K.K.) and visualized using NDP.view2 image viewing software (Hamamatsu Photonics K.K.). Necrosis percentage and mitosis counts were scored and averaged across 10 High-Power-Fields (HPF) at 40x magnification, then statistically tested (treatment vs. control using a one-sided, unpaired t-test).

### Cell viability assays

2–3×10^3^ EpS with/without DOX-induced *SMARCB1* re-expression were seeded in 90 µl medium per well (96-well). After 24 h, drugs were added to final concentration ranging from 0.01 µM to 50 µM with 0.05% dimethyl sulfoxide (DMSO) in all conditions. 48 h after the addition of the test drugs, cell viability was assessed using a Resazurin-based readout system^69^. Relative fluorescence units of treated wells were background corrected and normalized to vehicle controls.

### Statistical analysis

Data was analyzed in GraphPad PRISM 9 (GraphPad Software, San Diego, CA, USA). Where not otherwise specified, the statistical significance of differences between two experimental groups were tested using the two-tailed Wilcoxon Rank Sum / Mann-Whitney test with the Holm–Bonferroni method to account for multiple comparisons. *p*-values <0.05 were considered statistically significant. Data visualization was performed in Graphpad PRISM 9.

### Data availability

Original Affymetrix transcriptome profiling data have been deposited at the Gene Expression Omnibus (GEO) under the accession code GSE276634. Proteomics data has been deposited at the Proteomics Identifications Database (PRIDE) under PXD053945. ATAC-Seq and ChIP-Seq data are available from the corresponding author upon reasonable request.

## AUTHOR CONTRIBUTIONS

J.J. established isogenic cell line models, designed and performed functional *in vitro* and *in vivo* experiments including bioinformatic and histological analyses. F.C.A. performed *in vivo* experiments. F.F. carried out *in vitro* experiments and established isogenic cell line models. F.Z. assisted in *in vitro* experiments. M.C.G., S.O., A.B. and R.I. assisted in *in vivo* experiments and A.K.C. in histological analysis. R.W. assisted in the generation of cell line models. D.H. provided expertise in mass spectrometric proteome measurement. F.B. assisted in the generation of histological stains. T.G.P.G. designed and supervised the study, provided biological and technical guidance as well as laboratory infrastructure. All authors read and approved the final manuscript.

## ACKNOWLEDGEMENTS

We thank S. Kutschmann and N. Gmelin for excellent technical assistance. We thank the technicians of the DKFZ Core Facilities, German Cancer Research Center (DKFZ), Cellular Tools Core Facility, for assistance in the generation of stable isogenic EpS cell lines, the MS-Based Protein Analysis Core Facility for providing assistance with proteomic analysis, the NGS Core Facility for excellent sequencing services and the microarray unit for microarray processing. We thank the Core Facility for Light Microscopy, German Cancer Research Center (DKFZ), for the assistance in the processing of xenograft tissue slides.

---

**Supp. Fig. 1:**
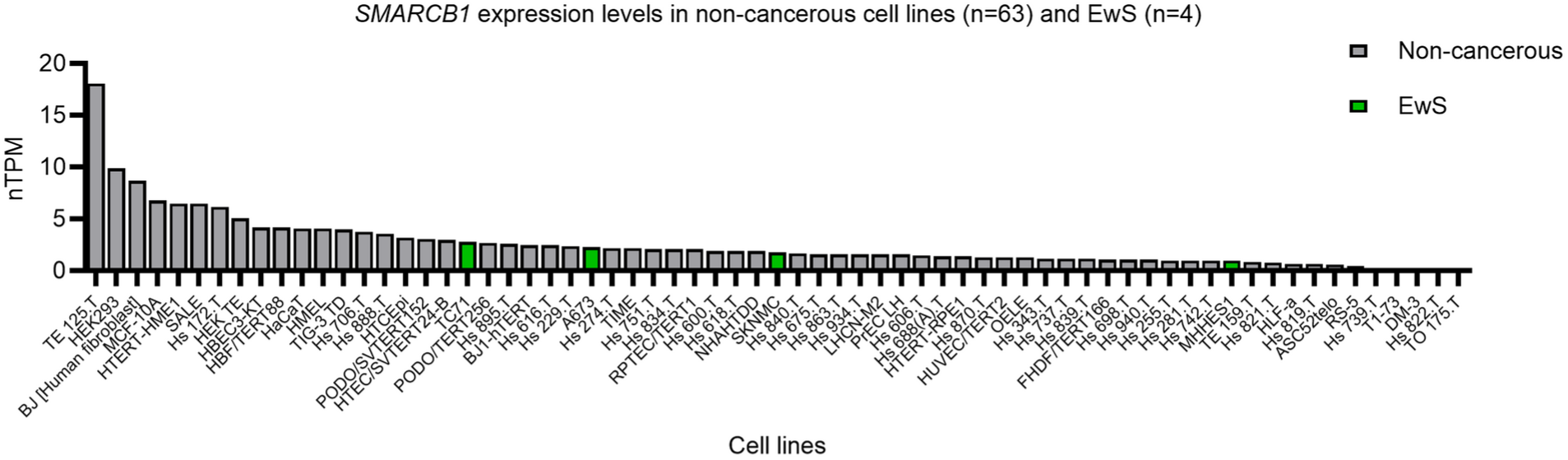
Expression levels (nTPM) of *SMARCB1* in non-cancerous cell lines (data source: Human Protein Atlas).

**Supp. Fig. 2:**
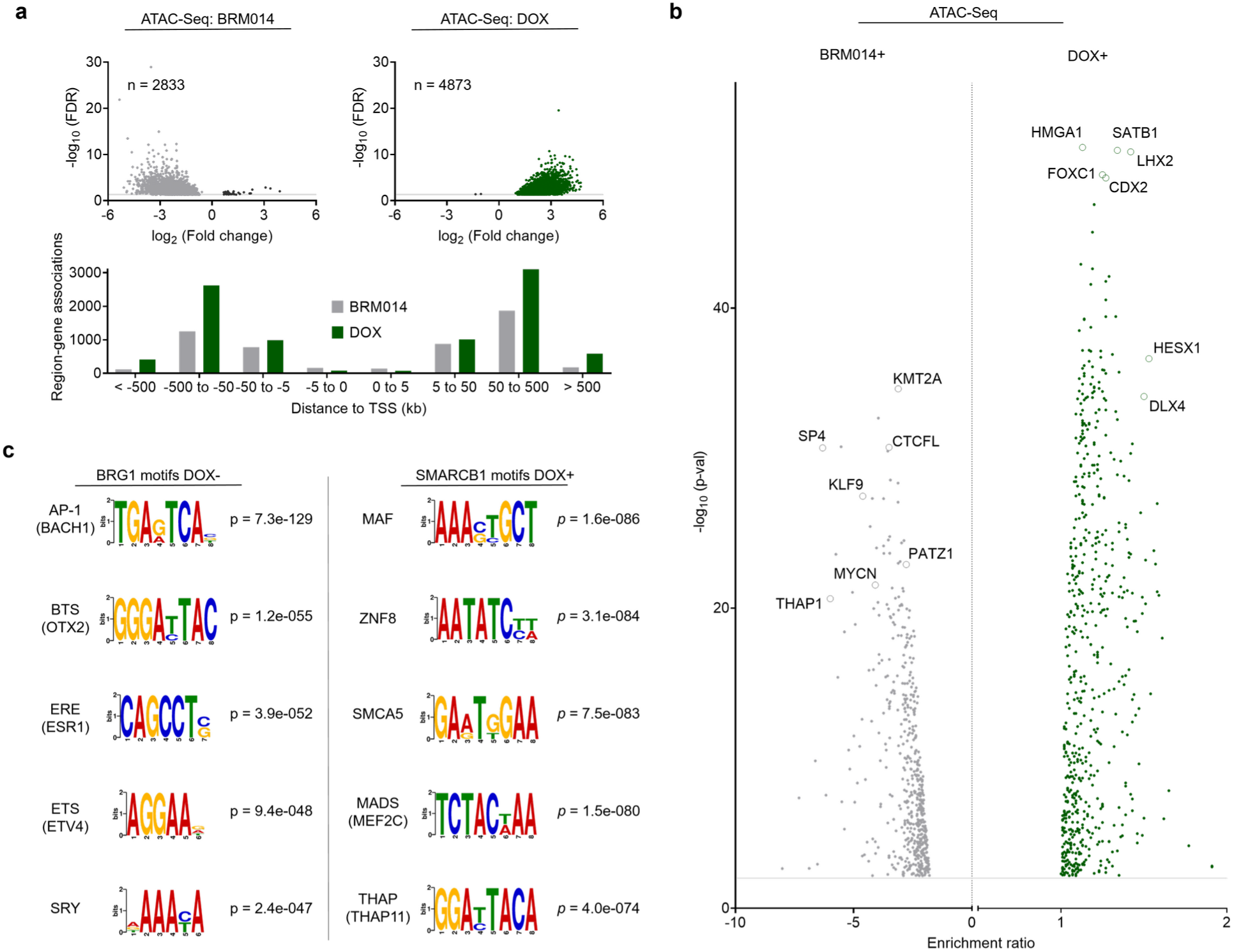
(a) Volcano plots of pooled differentially enriched open chromatin regions (FDR > 0.05) after BRM014 or DOX treatment alongside GREAT region-gene-associations binned by distance to closest gene TSS. (b) Volcano plot of Simple Enrichment Analysis (SEA) demonstrating highly enriched motifs in open chromatin regions lost upon BRM014 treatment and gained upon SMARCB1 re-expression (pooled across all EpS cell lines), each computed with the other condition as background sequences to show differential motif enrichment. (c) Top enriched gene motifs of BRG1 (DOX-) and INI-1 (DOX+) occupied loci in VA-ES-BJ.

**Supp. Fig. 3:**
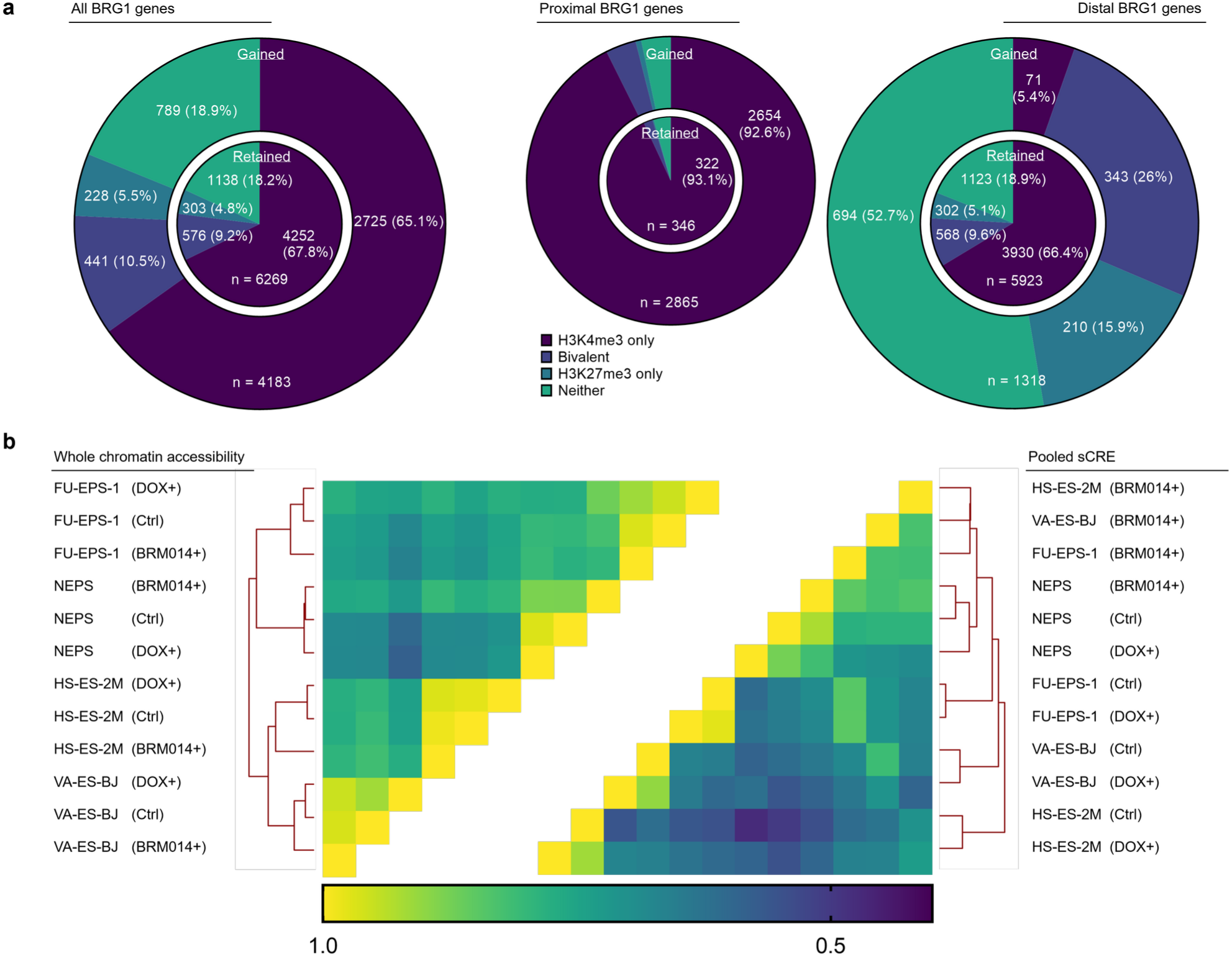
(a) Distribution of promoter categories of all, proximal (≤ 2kb from TSS) or distal (up to 1 Mb), retained (inner core) or gained (outer ring) GREAT associated BRG1 genes upon SMARCB1 re-expression. (b) Clustered Pearson correlation matrices of EpS cell line whole and differentially accessible chromatin regions.

**Supp. Fig. 4:**
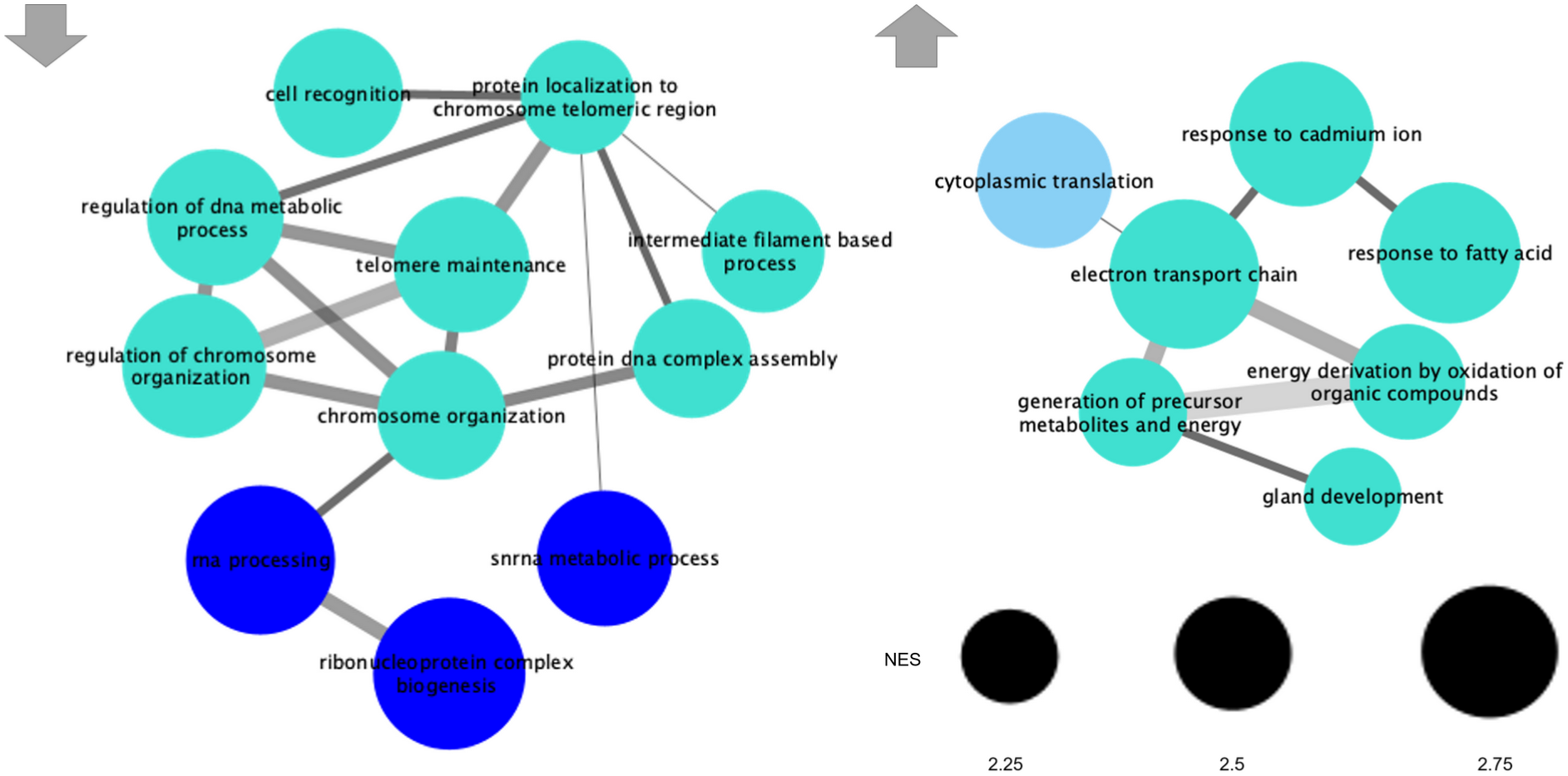
GSEA-based network analysis of up-/down-regulated biological process sets upon SMARCB1 re-expression from shared regulated proteins in VA-ES-BJ and NEPS after coIP against BRG1.

**Supp. Fig. 5:**
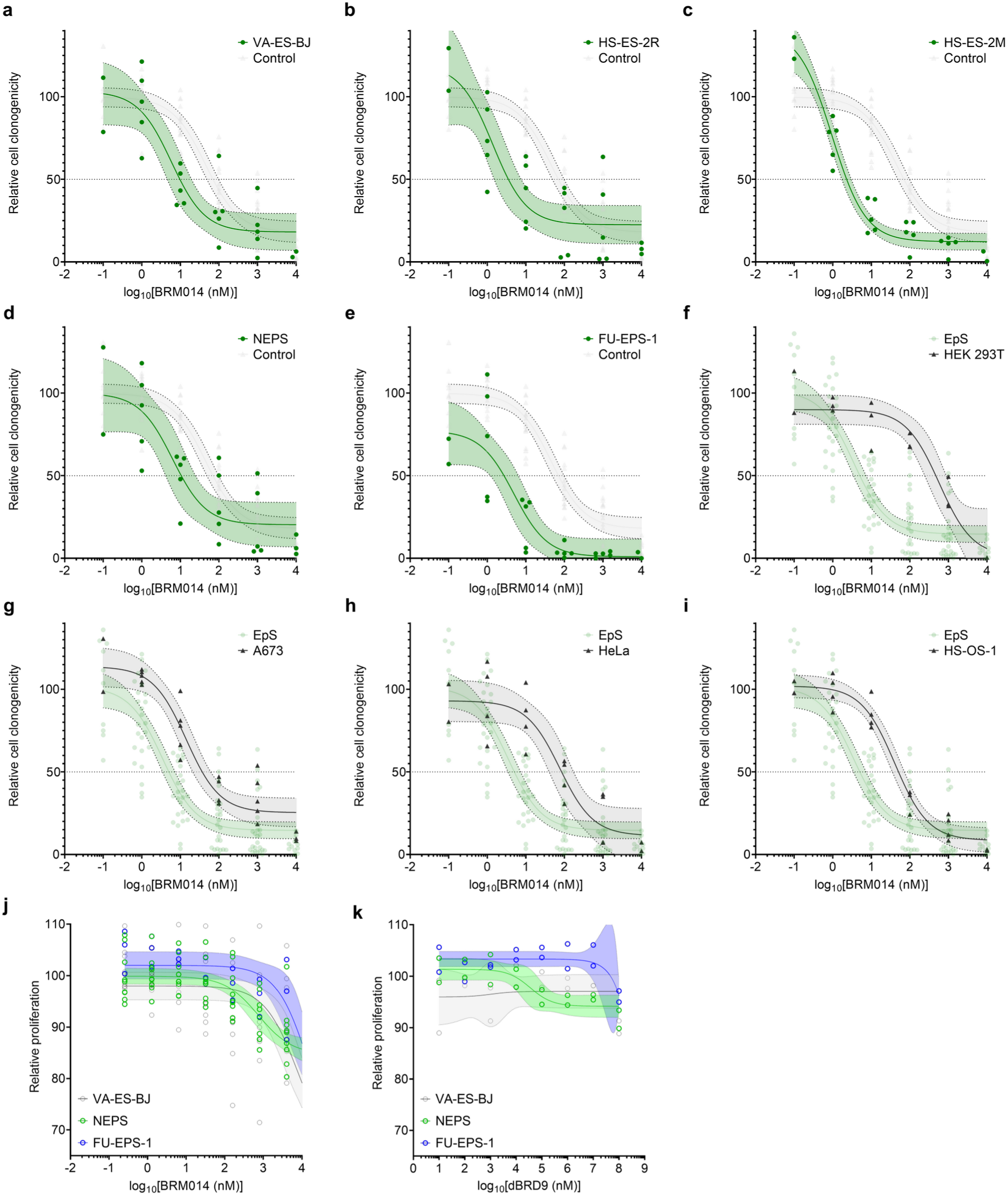
(a-i) Individual BRM014 drug response curves of SWI/SNF deficient EpS and control cell lines, calculated from clonogenicity (colony formation). (j) Resazurin metabolization based 48h BRM014 drug response with 95% CI of EpS cell lines. (k) Resazurin metabolization based 48h dBRD9 drug response with 95% CI of EpS cell lines.

**Supp. Fig. 6:**
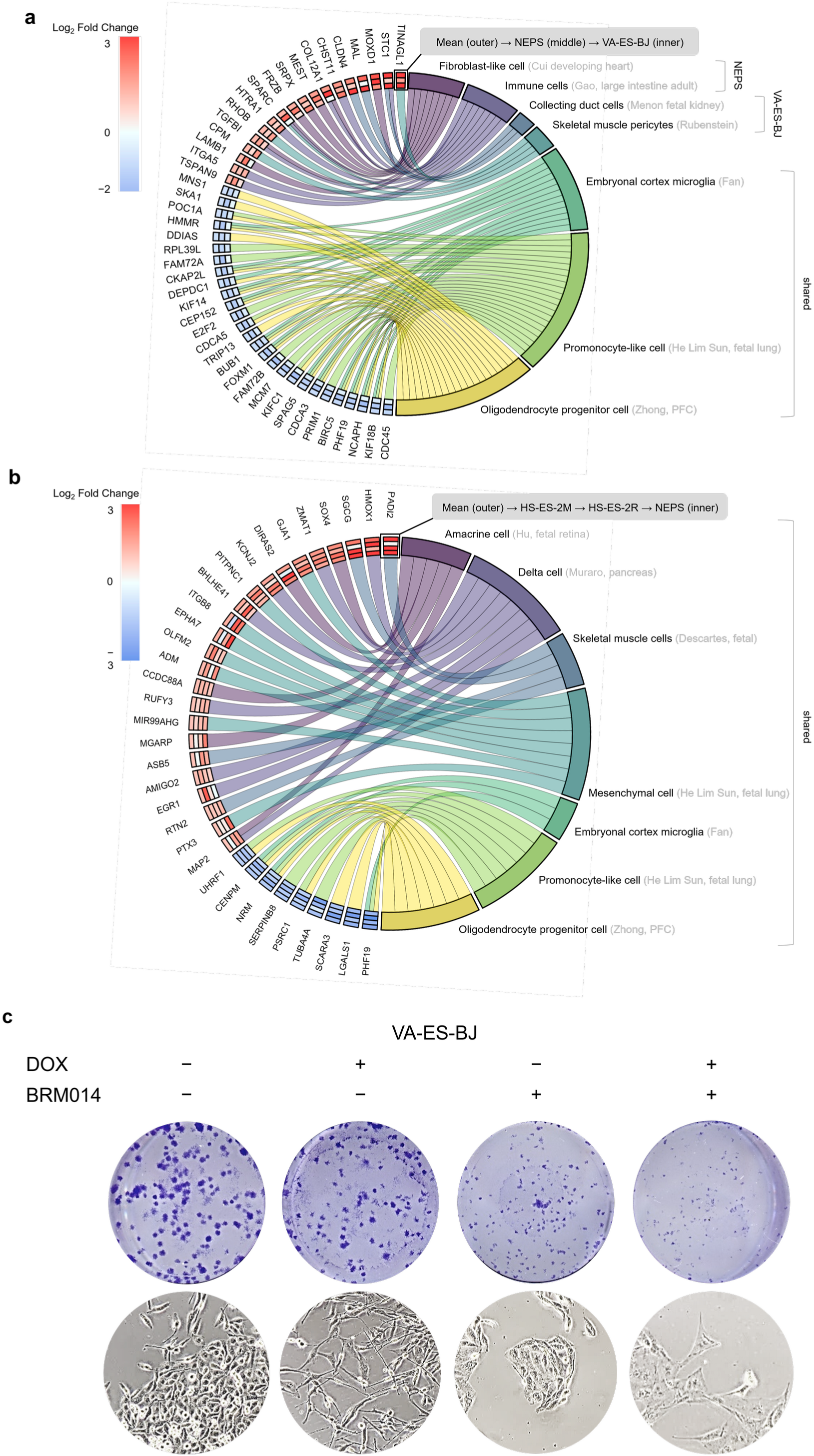
(a) Chord diagram showing the core genes of differentially regulated GSEA-based cell signatures for NEPS and VA-ES-BJ upon SMARCB1 re-expression as different annuli of log2 fold changes on the left (from outermost to innermost: Mean, NEPS, VA-ES-BJ) with cell signatures shown on the right. Left–right connections indicate gene membership in a signature’s leading-edge analysis. (b) Chord diagram showing the core genes of significantly regulated cell signatures for NEPS and VA-ES-BJ upon BRM014 treatment as different annuli of log2 fold changes on the left (from outermost to innermost: Mean, HS-ES-2M, HS-ES-2R, NEPS) with cell signatures shown on the right. Left–right connections indicate gene membership in a signature’s leading-edge analysis. (c) Colony formation assay and Brightfield images of VA-ES-BJ demonstrating synergistic action of SMARCB1 re-expression and SWI/SNF ATPase inhibition (via BRM014) as well as associated morphological changes.

## REFERENCES

1. Enzinger, F. M. Epithelioid sarcoma.A sarcoma simulating a granuloma or a carcinoma. Cancer 26, 1029–1041 (1970).

2. Noujaim, J. et al. Epithelioid Sarcoma: Opportunities for Biology-Driven Targeted Therapy. Frontiers in oncology 5, 186 (2015).

3. Casanova, M. et al. Epithelioid sarcoma in children and adolescents: a report from the Italian Soft Tissue Sarcoma Committee. Cancer 106, 708–717 (2006).

4. Jawad, M. U., Extein, J., Min, E. S. & Scully, S. P. Prognostic Factors for Survival in Patients with Epithelioid Sarcoma: 441 Cases from the SEER Database. Clin Orthop Relat Res 467, 2939–2948 (2009).

5. Armah, H. B. & Parwani, A. V. Epithelioid sarcoma. Arch Pathol Lab Med 133, 814–819 (2009).

6. Gounder, M. et al. Tazemetostat in advanced epithelioid sarcoma with loss of INI1/SMARCB1: an international, open-label, phase 2 basket study. The Lancet Oncology 21, 1423–1432 (2020).

7. Kazansky, Y. et al. Overcoming Clinical Resistance to EZH2 Inhibition Using Rational Epigenetic Combination Therapy. Cancer Discovery 14, 965–981 (2024).

8. Modena, P. et al. SMARCB1/INI1 Tumor Suppressor Gene Is Frequently Inactivated in Epithelioid Sarcomas. Cancer Research 65, 4012–4019 (2005).

9. Sullivan, L. M., Folpe, A. L., Pawel, B. R., Judkins, A. R. & Biegel, J. A. Epithelioid sarcoma is associated with a high percentage of SMARCB1 deletions. Modern pathology: an official journal of the United States and Canadian Academy of Pathology, Inc 26, 385–392 (2013).

10. Grünewald, T. G. P. et al. Translational Aspects of Epithelioid Sarcoma: Current Consensus. Clin Cancer Res 30, 1079–1092 (2024).

11. Kadoch, C. et al. Proteomic and Bioinformatic Analysis of mSWI/SNF (BAF) Complexes Reveals Extensive Roles in Human Malignancy. Nat Genet 45, 592–601 (2013).

12. Mashtalir, N. et al. A structural model of the endogenous human BAF complex informs disease mechanisms. Cell 183, 802–817.e24 (2020).

13. Mashtalir, N. et al. Modular Organization and Assembly of SWI/SNF Family Chromatin Remodeling Complexes. Cell 175, 1272–1288.e20 (2018).

14. Roberts, C. W. M., Leroux, M. M., Fleming, M. D. & Orkin, S. H. Highly penetrant, rapid tumorigenesis through conditional inversion of the tumor suppressor gene Snf5. Cancer Cell 2, 415–425 (2002).

15. Chatterjee, S. S., Biswas, M., Boila, L. D., Banerjee, D. & Sengupta, A. SMARCB1 Deficiency Integrates Epigenetic Signals to Oncogenic Gene Expression Program Maintenance in Human Acute Myeloid Leukemia. Molecular Cancer Research 16, 791–804 (2018).

16. Zhu, Z. et al. Mitotic bookmarking by SWI/SNF subunits. Nature (2023) doi:10.1038/s41586-023-06085-6.

17. Custers, L. et al. Somatic mutations and single-cell transcriptomes reveal the root of malignant rhabdoid tumours. Nat Commun 12, 1407 (2021).

18. Betz, B. L., Strobeck, M. W., Reisman, D. N., Knudsen, E. S. & Weissman, B. E. Re-expression of hSNF5/INI1/BAF47 in pediatric tumor cells leads to G1arrest associated with induction of p16ink4a and activation of RB. Oncogene 21, 5193–5203 (2002).

19. Jagani, Z. et al. Loss of the tumor suppressor Snf5 leads to aberrant activation of the Hedgehog-Gli pathway. Nat Med 16, 1429–1433 (2010).

20. Mora-Blanco, E. L. et al. Activation of β-catenin/TCF targets following loss of the tumor suppressor SNF5. Oncogene 33, 933–938 (2014).

21. Wolf, B. K. et al. Cooperation of chromatin remodeling SWI/SNF complex and pioneer factor AP-1 shapes 3D enhancer landscapes. Nat Struct Mol Biol 30, 10–21 (2023).

22. Mashtalir, N. et al. Chromatin landscape signals differentially dictate the activities of mSWI/SNF family complexes. Science 373, 306–315 (2021).

23. Nakayama, R. T. et al. SMARCB1 is required for widespread BAF complex-mediated activation of enhancers and bivalent promoters. Nat Genet 49, 1613–1623 (2017).

24. Brenca, M. et al. SMARCB1/INI1 genetic inactivation is responsible for tumorigenic properties of epithelioid sarcoma cell line VAESBJ. Molecular cancer therapeutics 12, 1060–1072 (2013).

25. Wang, X. et al. SMARCB1-mediated SWI/SNF complex function is essential for enhancer regulation. Nat Genet 49, 289–295 (2017).

26. Rago, F. et al. Exquisite Sensitivity to Dual BRG1/BRM ATPase Inhibitors Reveals Broad SWI/SNF Dependencies in Acute Myeloid Leukemia. Molecular Cancer Research 20, 361–372 (2022).

27. Xiao, L. et al. Targeting SWI/SNF ATPases in enhancer-addicted prostate cancer. Nature 601, 434–439 (2022).

28. Wang, X. et al. Oncogenesis caused by loss of the SNF5 tumor suppressor is dependent on activity of BRG1, the ATPase of the SWI/SNF chromatin remodeling complex. Cancer Res 69, 8094–8101 (2009).

29. Moe, K. C. et al. The SWI/SNF ATPase BRG1 facilitates multiple pro-tumorigenic gene expression programs in SMARCB1-deficient cancer cells. Oncogenesis 11, 30 (2022).

30. Orth, M. F. et al. Systematic multi-omics cell line profiling uncovers principles of Ewing sarcoma fusion oncogene-mediated gene regulation. Cell Rep 41, 111761 (2022).

31. Uhlen, M. et al. A pathology atlas of the human cancer transcriptome. Science 357, eaan2507 (2017).

32. Franken, N. A. P., Rodermond, H. M., Stap, J., Haveman, J. & van Bree, C. Clonogenic assay of cells in vitro. Nat Protoc 1, 2315–2319 (2006).

33. Papillon, J. P. N. et al. Discovery of Orally Active Inhibitors of Brahma Homolog (BRM)/SMARCA2 ATPase Activity for the Treatment of Brahma Related Gene 1 (BRG1)/SMARCA4-Mutant Cancers. Journal of medicinal chemistry 61, 10155–10172 (2018).

34. Radko-Juettner, S. et al. Targeting DCAF5 suppresses SMARCB1-mutant cancer by stabilizing SWI/SNF. Nature 628, 442–449 (2024).

35. Schick, S. et al. Acute BAF perturbation causes immediate changes in chromatin accessibility. Nat Genet 53, 269–278 (2021).

36. Iurlaro, M. et al. Mammalian SWI/SNF continuously restores local accessibility to chromatin. Nat Genet 53, 279–287 (2021).

37. Chen, E. Y. et al. Enrichr: interactive and collaborative HTML5 gene list enrichment analysis tool. BMC Bioinformatics 14, 128 (2013).

38. Kuleshov, M. V. et al. Enrichr: a comprehensive gene set enrichment analysis web server 2016 update. Nucleic Acids Res 44, W90–97 (2016).

39. Fujioka, S. et al. NF-κB and AP-1 Connection: Mechanism of NF-κB-Dependent Regulation of AP-1 Activity. Mol Cell Biol 24, 7806–7819 (2004).

40. Remillard, D. et al. Degradation of the BAF Complex Factor BRD9 by Heterobifunctional Ligands. Angew Chem Int Ed Engl 56, 5738–5743 (2017).

41. Wang, X. et al. BRD9 defines a SWI/SNF sub-complex and constitutes a specific vulnerability in malignant rhabdoid tumors. Nat Commun 10, 1881 (2019).

42. Michel, B. C. et al. A non-canonical SWI/SNF complex is a synthetic lethal target in cancers driven by BAF complex perturbation. Nat Cell Biol 20, 1410–1420 (2018).

43. Brien, G. L. et al. Targeted degradation of BRD9 reverses oncogenic gene expression in synovial sarcoma. eLife https://elifesciences.org/articles/41305 (2018) doi:10.7554/eLife.41305.

44. Kurata, K. et al. BRD9 Degradation Disrupts Ribosome Biogenesis in Multiple Myeloma. Clin Cancer Res 29, 1807–1821 (2023).

45. Panditharatna, E. et al. BAF Complex Maintains Glioma Stem Cells in Pediatric H3K27M Glioma. Cancer discovery 12, 2880–2905 (2022).

46. Mo, Y. et al. Epigenome programing by H3.3K27M mutation creates a dependence of pediatric glioma on SMARCA4. Cancer Discov 12, 2906–2929 (2022).

47. Harttrampf, A. C., da Costa, M. E. M., Renoult, A., Daudigeos-Dubus, E. & Geoerger, B. Histone deacetylase inhibitor panobinostat induces antitumor activity in epithelioid sarcoma and rhabdoid tumor by growth factor receptor modulation. BMC Cancer 21, 833 (2021).

48. Lopez, G. et al. HDAC Inhibition for the Treatment of Epithelioid Sarcoma: Novel Cross Talk Between Epigenetic Components. Mol Cancer Res 14, 35–43 (2016).

49. Fiskus, W. et al. BRG1/BRM inhibitor targets AML stem cells and exerts superior preclinical efficacy combined with BET or menin inhibitor. Blood 143, 2059–2072 (2024).

50. DiNardo, C. D. et al. Preliminary Results from a Phase 1 Dose Escalation Study of FHD-286, a Novel BRG1/BRM (SMARCA4/SMARCA2) Inhibitor, Administered As an Oral Monotherapy in Patients with Advanced Hematologic Malignancies. Blood 142, 4284 (2023).

51. Akhurst, R. J. & Derynck, R. TGF-beta signaling in cancer--a double-edged sword. Trends Cell Biol 11, S44–51 (2001).

52. Massagué, J. & Sheppard, D. TGF-β signaling in health and disease. Cell 186, 4007–4037 (2023).

53. Karlsson, M. et al. A single-cell type transcriptomics map of human tissues. Sci Adv 7, eabh2169 (2021).

54. Marchetto, A. & Romero-Pérez, L. Western Blot Analysis in Ewing Sarcoma. Methods Mol Biol 2226, 15–25 (2021).

55. Marchetto, A. et al. Oncogenic hijacking of a developmental transcription factor evokes vulnerability toward oxidative stress in Ewing sarcoma. Nat Commun 11, 2423 (2020).

56. Subramanian, A. et al. Gene set enrichment analysis: A knowledge-based approach for interpreting genome-wide expression profiles. Proceedings of the National Academy of Sciences 102, 15545–15550 (2005).

57. Langfelder, P. & Horvath, S. WGCNA: an R package for weighted correlation network analysis. BMC Bioinformatics 9, 559 (2008).

58. Waszak, S. M. et al. Germline Elongator mutations in Sonic Hedgehog medulloblastoma. Nature 580, 396–401 (2020).

59. Walter, W., Sánchez-Cabo, F. & Ricote, M. GOplot: an R package for visually combining expression data with functional analysis. Bioinformatics 31, 2912–2914 (2015).

60. Shevchenko, A., Tomas, H., Havli, J., Olsen, J. V. & Mann, M. In-gel digestion for mass spectrometric characterization of proteins and proteomes. Nat Protoc 1, 2856–2860 (2006).

61. Tyanova, S., Temu, T. & Cox, J. The MaxQuant computational platform for mass spectrometry-based shotgun proteomics. Nat Protoc 11, 2301–2319 (2016).

62. Cox, J. et al. Accurate Proteome-wide Label-free Quantification by Delayed Normalization and Maximal Peptide Ratio Extraction, Termed MaxLFQ. Mol Cell Proteomics 13, 2513–2526 (2014).

63. Schwanhäusser, B. et al. Global quantification of mammalian gene expression control. Nature 473, 337–342 (2011).

64. Wieczorek, S. et al. DAPAR & ProStaR: software to perform statistical analyses in quantitative discovery proteomics. Bioinformatics 33, 135–136 (2017).

65. Stark, R. & Brown, G. DiffBind. Bioconductor http://bioconductor.org/packages/DiffBind/ (2011).

66. McLean, C. Y. et al. GREAT improves functional interpretation of cis-regulatory regions. Nat Biotechnol 28, 495–501 (2010).

67. Tanigawa, Y., Dyer, E. S. & Bejerano, G. WhichTF is functionally important in your open chromatin data? PLoS Comput Biol 18, e1010378 (2022).

68. Enot, D. P., Vacchelli, E., Jacquelot, N., Zitvogel, L. & Kroemer, G. TumGrowth: An open-access web tool for the statistical analysis of tumor growth curves. Oncoimmunology 7, (2018).

69. Musa, J. & Cidre-Aranaz, F. Drug Screening by Resazurin Colorimetry in Ewing Sarcoma. in Ewing Sarcoma: Methods and Protocols (eds. Cidre-Aranaz, F. & G. P. Grünewald, T.) 159–166 (Springer US, New York, NY, 2021). doi:10.1007/978-1-0716-1020-6_12.

